# Mapping of genotype-by-environment interaction loci for Metabolic Syndrome-like traits using the multi-parent *Drosophila* Synthetic Population Resource determines that main genetic effects are distinct from environment dependent plastic loci

**DOI:** 10.1101/2025.08.04.668530

**Authors:** Kelly Dew-Budd, Ravi Mathur, Siddharth Roy, Julie Jarnigan, Andrea Moss, Andrei Bombin, Vishal Oza, Chinmay Rele, Alison Adams, Sean Mendez, Katherine Bray, Dana Davis, Matthew Kieffer, Leah Leonard, Joana Hubickey, Cheyenne Paiva, Nicholas Izor, Divya Nadella, Lauren Ross Perkins, Xiangpei Zeng, Jordyn Merriam, Alison Motsinger-Reif, Laura K. Reed

## Abstract

Metabolic Syndrome (MetS) risk, driven by genotype-environment interactions like diet, is rising globally. Due to its genetic and environmental complexity, the genetic architecture and interconnected traits underlying MetS is poorly understood. In *Drosophila*, genotype-by-diet interactions significantly influence MetS-like traits. This study used the *Drosophila* Synthetic Population Resource to dissect the genetic architecture of both genotypic and genotype-by-diet interaction effects underlying trait variation. The study hypotheses were: 1) Loci responsible for metabolic phenotypic variation should be shared across traits. 2) Genetic loci responsible for plasticity and epistatic interactions for metabolic traits should also be the loci responsible for the main effects. 3) Genes responsible for variation in metabolic traits should share common functions. Using a round-robin crossing scheme and novel analyses, we mapped additive, dominance, and epistatic loci—some diet-specific, others diet-independent. Main-effect and plastic loci were largely distinct, as were epistatic loci from main-effect loci, highlighting that main genetic effects alone will not explain how genetic variants interact with the environment or the genome to influence disease risk. gene-by-diet or gene-by-gene interactions influencing MetS risk. Further, tremendous cryptic genetic variation for metabolic traits is lurking in natural populations. We explored the function of candidate genes from our study empirically and with bioinformatics. While some of the candidate genes might have been expected, most would not have been identified *a priori*, thus with this study we have identified many new candidate mechanisms contributing to the genetic and genotype-by-diet interaction effects on MetS variance.

## INTRODUCTION

A rapidly growing imbalance between energy intake and expenditure has resulted in increasing rates of Metabolic Syndrome (MetS) (Basciano *et al*. 2005; Alberti *et al*. 2006; Kaur 2014; Hulsegge *et al*. 2016). This disease is characterized by symptoms such as insulin resistance, abdominal obesity, and dyslipidemia (Alberti *et al*. 2006). The epidemic proportions of MetS, as well as its high correlation with the development of type-2 diabetes and cardiovascular disease, increases the importance of research into its causes and potential clinical prevention strategies (Basciano *et al*. 2005). MetS is composed of complex quantitative traits, influenced by interactions between genetic and environmental factors (O’Rahilly and Farooqi 2006; Farooqi and O’Rahilly 2006). Our understanding of these quantitative traits is essential for translation of genomic findings into clinical practice (Corella *et al*. 2009). Therefore, elucidating the genetic components of these complex quantitative traits is vital to understanding MetS as a whole.

*Drosophila melanogaster* can be used to model MetS, as many of their metabolic pathways are homologous to human pathways (Trinh and Boulianne 2013). These metabolic pathways include the storage and utilization of energy, signaling from the central nervous system to regulate food intake and satiation, sugar homeostasis, and lipid homeostasis (Trinh and Boulianne 2013). An example of this homology is shown when insulin-producing cells of *Drosophila* are ablated the flies exhibit a phenotype comparable to type-1 diabetes (Broughton *et al*. 2005; Rulifson *et al*. 2002). In this study, we used body weight, triglyceride storage, and sugar content as phenotypes representing similarly metabolically controlled traits in humans (body mass index, adiposity, and blood sugar), thus we refer to these as MetS-like traits.

While the genetic effect on a phenotype can be tested by measuring a quantitative trait in multiple genotypes, the genotype-by-environment interaction effect is determined by testing each genotype in more than one environment. An interaction occurs when the difference between the measurements for the two environments is not equal for every genotype, indicating the subjects reacted differently to the distinct environments. Genetic mapping of MetS-related traits in humans has identified thousands of loci contributing to these traits across populations (e.g., 4277 associations for type-2 diabetes mellitus (EFO_0001360) downloaded 8/16/22 from the NHGRI-EBI GWAS Catalog (Buniello, MacArthur *et al*. 2019)), thus they are clearly highly polygenic. Further, statistical epistatic interactions contributing to genetic variance in MetS-related traits in humans and other model organisms have also been observed (e.g., De Luca *et al*. 2005, Yi *et al*. 2004, De *et al*. 2015, Smemo *et al*. 2014).

Historically, it has been assumed that the genetic loci responsible for the additive genetic variance in a population are also the ones responsible for any genetic interactions with the environment (plasticity) and other genetic loci (epistasis). A major reason for this assumption was driven by limitations on statistical and computational power in mapping studies (e.g., the statistical tests for pairwise genetic interactions scale as the square of the number of loci; Mackay, 2014), thus traditionally, interactions were only tested for if a main effect was first identified. Further, additive models generally do a reasonable job of describing the majority overall heritable variance for a quantitative trait in a population, leading many to erroneously assume that genetic interactions are unimportant at the level of the locus (Hill *et al*. 2008, Crow 2010, Huang and Mackay 2016). However, growing numbers of studies demonstrate that when considered from the level of the individual genetic locus, much of the apparent heritable genetic variance at the population levels is in fact driven by interactions (Lehner *et al*. 2006, Ooi *et al*. 2006, Huang *et al*. 2012, Huang and Mackay 2016, Mackay 2014, Tyler *et al*. 2016, Forsberg *et al*. 2017).

Our previous work has quantified the relative contribution of genotype, diet, and genotype-by-diet interactions to variation in MetS-like traits in *Drosophila* (Reed *et al*. 2010; 2014; Williams *et al*. 2015), finding that the genotype-by-diet interaction effects contribute substantially to the population phenotypic variance. Motivated by that finding, and by work in other systems, in this study we used genetic mapping to identify loci responsible for the genotype-by-environment interactions both for main and epistatic effects. Figure 1 provides a visual example of the genotype-phenotype relationship for genetic and genotype-by-environment interacting loci for main and epistatic effects. We compared the phenotypic response to a normal and high fat diet to reflect the change in diet of westernized cultures, which may be contributing to the rise of MetS (O’Rahilly and Farooqi 2006). The primary questions we endeavored to answer were: 1) Are common loci responsible for phenotypic variation across MetS-like traits? 2) Are genetic loci of main effect also those responsible for genetic variation in environmental plasticity and epistatic interactions for MetS-like traits? 3) If loci largely do differ across traits and interaction effects, are there any core genes or gene functions that are shared?

**Figure 1:**
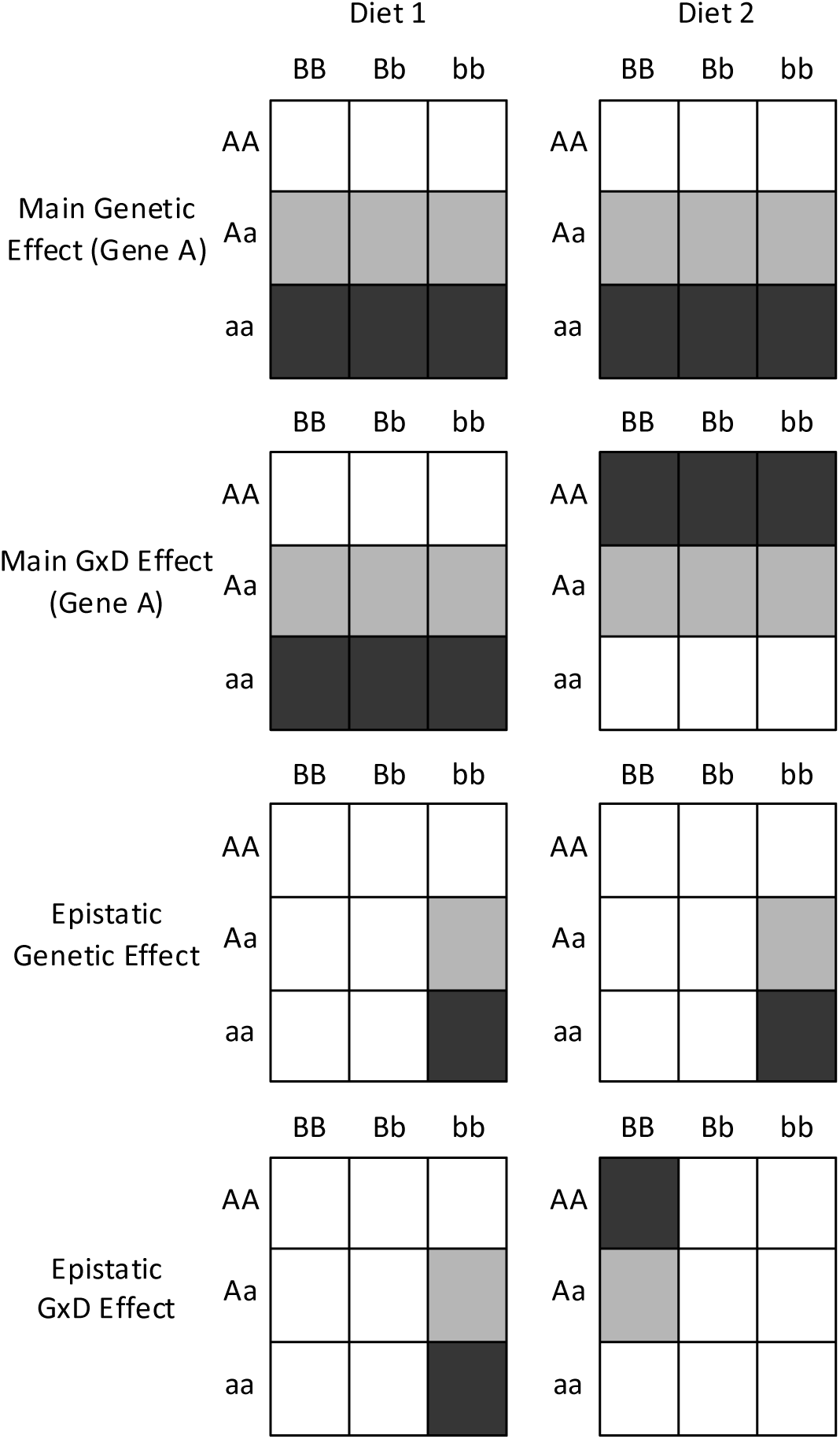
Example of relationship between genotype and phenotype due to genetic, genotype-by-diet, epistatic, and epistasis by diet QTLs. The scenarios include two possible genetic loci (A and B) and two environments (Diets 1 and 2). Phenotype is indicated by the degree of shading of a box in the matrix, and each box represents a different genotype at the two genetic loci. The top two panels depict a main genetic effect QTL at locus A where the phenotype is influenced by the genotype of A in the same way on both diets. The second row of panels depicts a genotype-by-diet interaction QTL at Locus A, where the phenotype varies with the genotype at A but in opposing directions on the two different diets. The third row depicts main epistatic effect QTLs, where the phenotype does not vary with diet but is dependent on the genotypes of both the A and B locus. Phenotypic variation due to A is only observed when the B locus has the *bb* genotype. The bottom row is an example of a genotype-by-diet epistatic effect, where, as above, the phenotype is determined by an interaction between the A and B locus, however the specific phenotype of that interaction differs between the two diets. There are many other genotype-by-phenotype relationships that would be consistent with interactions between genotype and diet and/or involving epistatic interactions, those presented here are just examples.

In this study, we used Population A RILs (recombinant inbred lines) of the *Drosophila* Synthetic Population Resource (DSPR) (King *et al*. 2012a; b), employing a round-robin design, to reestablish heterozygous genotypes. Another study within this population identified substantial genotype-by-diet interaction effects on lifespan and fecundity as evidenced by low genetic (Ng’oma *et al*, 2018). We identified influential loci for genotype-by-diet interactions and epistatic effects contributing to male and female pupal body weight and larval triglyceride and trehalose concentration.

We determined that the loci responsible for the main genetic effects on these traits were distinct from those observed for the genotype-by-diet interactions, for both the non-epistatic and epistatic loci. Due to the round-robin design, we were able to identify additive, dominance, and full founder-by-parent genetic effects for the non-epistatic loci. The maximum phenotypic variance explained by a significant locus for any trait was 5.1% for triglycerides, while most of the significant loci only explained a few percent of the total phenotypic variation. In addition, we found little overlap between the non-epistatic loci across MetS-like traits, which suggests that the generally observed correlations among these traits are driven primarily by very small effect loci. Gene ontology enrichment of the genes associated with the quantitative trait loci (QTLs) identified both familiar and novel potential mechanisms underlying the phenotypic variation. Candidate genes from four of the genotype-by-diet QTL were selected for functional experiments. Using both gene expression and mutant analysis, we were able to identify genes that may be strong candidates for causing a genotype-by-diet interaction in *Drosophila*.

## MATERIALS AND METHODS

### QTL Discovery

#### Drosophila Stocks

We used 604 RILs from Population A of the *Drosophila* Synthetic Population Resource (DSPR) obtained from S. Macdonald for QTL discovery (King *et al*. 2012a; b). All stocks were maintained at 25°C at 50% humidity on a 12 hr light-dark cycle and a standard cornmeal-molasses diet (defined as “normal” in Reed *et al*. 2010) for two or more generations prior to experimentation. The DSPR was originally generated under constant 24-hour light conditions (King *et al*. 2012a), so there may have been some selection for modified circadian rhythms in the creation of the population. However, since circadian light:dark cycles are important for metabolic homeostasis (e.g., Bray and Young 2007; Laposky *et al*. 2008) in this study we used a 12 hr light-dark cycle to better emulate natural conditions. It is possible that a similar mapping study performed with no light cycle would map different loci than those mapped here. For experimentation, flies were housed in the same controlled environment, but subsets of each genotype were raised on either the normal or high fat diet (Figure 2). The high fat diet consisted of the normal diet supplemented with 3% coconut oil by weight, which resulted in an increase in saturated fat by 0.26 g per 10 g vial of food (Reed *et al*. 2010; *USDA National Nutrient Database for Standard Reference* 2015).

**Figure 2:**
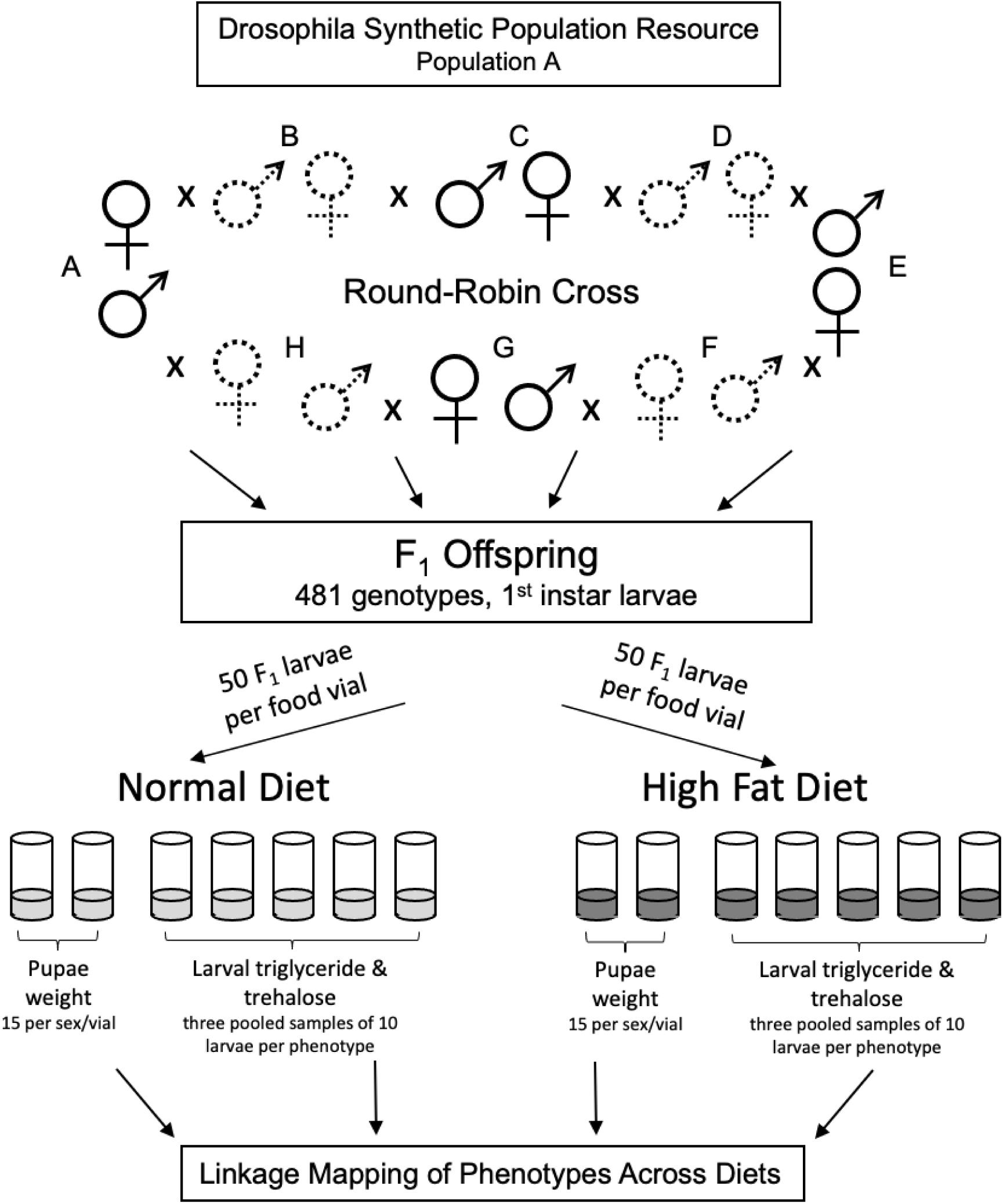
Experimental design for generating fly samples. Within one round of round-robin crosses, conducted simultaneously, females of genotype A were crossed to males of genotype B, and females from genotype B were crossed to males of genotype C etc. The F_1_ offspring of the crosses were raised at a density of 50 larvae per food vial and then collected for phenotype measurements. Two food vials per diet and genotype were used to collect pupae for weighing (15 pupae per sex per vial) and five food vials per diet were pooled to collect pooled third instar larvae for triglyceride and trehalose measurements (ten larvae per sample, three samples per phenotype). These phenotype measurements were then genetically mapped.

#### Experimental Design and Procedures

Virgins were collected using CO_2_ and were then housed separately for 3-4 days to confirm virgin status. Using a round-robin crossing scheme (females from one line mated to males of the line in sequential order) with 20-30 lines per round, 30 virgin females and 15 males were placed in laying chambers with an apple agar plate supplemented with reconstituted active dry yeast. Agar plates were changed every 12 hours and first instar larvae were collected 24 hours after eggs were laid. Larvae were placed on control and high fat diets at a density of 50 larvae per vial with 10g of food. Third-instar larvae were collected 72 hours later and fasted on a water agar plate for 3-4 hours. Samples of thirty larvae were frozen in liquid nitrogen and stored at -80°C for RNA expression analysis and replicates of ten larvae were collected and stored at -20°C for triglyceride and trehalose measurements, up to three replicates per assay. Triglyceride levels were determined from homogenized larvae with the Sigma-Aldrich Triglyceride Determination Kit (Clark and Keith 1988; De Luca *et al*. 2005; Reed *et al*. 2010) and were used as a corollary to adiposity variation in metabolic disorders.

Trehalose levels were tested using the Sigma-Aldrich Glucose (GO) assay kit following overnight digestion with trehalase of larval homogenates as in previous studies (Reed *et al*. 2014; Williams *et al*. 2015). Trehalose levels were used to measure hemolymph sugar content as a corollary to variation in blood sugar found in metabolic disorders. Two technical replicates were used for each sample and 2-3 samples were tested for each cross on each diet. Two additional food vials of larvae for each genotype-by-diet treatment combination were allowed to pupate and were collected ∼12 hours before eclosion (when the male sex combs could be visualized through the pupal case), and frozen in Ringer’s solution at -20°C. Stored pupae were cleaned and sorted by sex, then weighed individually (wet weight). Up to 15 pupae of each sex were weighed for each food vial (Reed *et al*. 2010). Weight is the corollary to body mass index and variation growth patterns found in metabolic disorders.

#### QTL Mapping Statistical Analysis

All mapping analysis was performed in R (R Development Core Team version 3.2.2 2014) and code to execute the analysis can be found at the Github site (https://github.com/rmathur87/DLinkMaP/tree/master). Founder haplotype probabilities for the DSPR RILs were obtained from the DSPRqtl and DSPRqtlDataA R packages available at http://FlyRILs.org/. The DSPR resource reports eight probabilities for each RIL at 10kB intervals, which corresponds to the probability that a RIL shares its haplotype with each founder. Nearly the entire genome for each RIL is homozygous and King *et al*. (King *et al*. 2012a) reports a high degree of certainty in these founder assignments.

We performed a mixed model analysis (detailed in the Supplemental Methods, File S1) at each location to test for linkage. Random effect terms corresponding to month, round robin group, RIL by month, and cross were included as both average effects over normal and high fat diet along with diet interaction terms. We also included a term for vial, which results in nine random effect groups in the initial model. We did not include terms corresponding to maternal or paternal specific line or reciprocal terms because these were not identifiable due to the round robin design. On the other hand, we found we could compute and test founder probabilities corresponding to all 8 X 8 = 64 founder-by-maternal/paternal combinations. We thus were able to fit six separate non-null models corresponding to additive, dominant, and full founder-by-parent effects as main effects and diet interactions. Statistical inference was conducted by calculating the χ2-statistics based on the difference in model log-likelihoods in a hierarchical manner, and determining p-values based on the χ2-distribution for additive, dominant, full, main (full compared to null), additive-by-diet, dominant-by-diet, full-by-diet, and diet (full-by-diet compared to full) effects.

The epistatic effects were incorporated by the cross product of the haplotype frequencies for the additive model only. Both main effects and all diet interactions were also included in the model along with all previously listed random effects. As in the non-epistatic analysis, six non-null models were fitted, all based on the additive model (single QTL, additive QTLs, main epistatic interaction, single QTL-by-diet, additive QTLs by diet, and diet epistatic interaction). Complete details of the epistatic model are included in the Supplemental Methods (File S1).

Epistatic effects for all pairwise 10kB positions in the genome were analyzed. As in the non-epistatic analysis, the statistical inference is based on χ2-statistics and negative log p-values are reported. The inference was conducted in a stepwise manner in the same way as the non-epistatic analysis. The significance threshold was set at a negative log p-value greater than six (NLP > 6), being more conservative than a Bonferroni correction. False discovery rates and QQ plots were also generated for the non-epistatic mapping results (Supplemental Methods, File S1).

For each significant peak, the 95% confidence intervals were computed using a LOD drop of two, as has been shown to be effective in previous work (Manichaikul *et al*. 2006; Broman and Sen 2009; King *et al*. 2014). To verify the significance of the peaks, permutation testing was performed (1,000 permutations) for the male weight phenotype (Supplemental Methods, File S1). For the epistatic models, a marginal confidence interval was calculated for each QTL involved in a significant epistatic interaction as determined by the threshold of six for the negative log p-value (Supplemental Methods, File S1). A Bayesian model was conducted to estimate the variance explained for each non-epistatic peak and the significant epistatic interactions (Supplemental Methods, File S1). Due to the non-independence of adjacent epistatic loci in the pairwise-interaction analysis approach, loci were clustered by genomic position for each if the loci were less than 0.35 kb from each other. Using each group as a node, interaction networks were generated by drawing an edge between two groups if at least one significant peak was common between the groups.

To perform gene expression analysis for candidate genes associated with genotype-by-diet interaction QTLs for triglycerides, we computed diet effects for each cross using only the haplotype frequencies for the model that identified the peak. This allowed us to prioritize a subset of the most informative cross-by-diet interactions for qPCR experiments. Additionally, the normalized triglyceride values were analyzed by ANOVA according to the model:

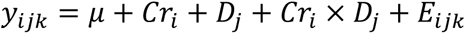

where Cr is the cross effect, D is the diet effect, Cr x D is the interaction term between cross and diet, and E is the error associated with the ith cross, jth diet, and kth individual.

### Candidate Gene Ontology Enrichment Analysis

Given the large number of QTL peaks identified across these mapping analyses, we identified candidate genes for each QTL and performed functional gene ontology enrichment analysis (Supplemental Methods, File S1) to facilitate biological interpretation and prioritize genes or pathways for functional validation. We performed functional enrichments analyses using G:profiler g:GOSt (Gene Group Functional Profiling version r1705_e86_eg33, Reimand *et al*. 2016), to test for overrepresentation of functional categories within three groups to prioritize genes for functional follow up. The three groups were: 1) genes that are associated with a phenotype due to genetic effects only, 2) genes that are associated with a phenotype due to the diet effects in a genetically specific way, and 3) all genes that are associated with each phenotype. Functional enrichment was described “significant” with a nominal p-value cut-off of 0.05; however, categories with greater significance were given preference in biological interpretation. To take advantage of the more extensive functional annotation of human orthologs, we also used bioDBnet (Mudunuri *et al*. 2009) to convert *Drosophila* genes’ IDs to known human homologous genes and repeated gene ontology enrichment analysis with the *Homo sapiens* annotation (Supplemental Methods, File S1).

The lists of candidate genes were also systematically compared to lists of genes generated in other studies of related traits from expression profiling, RNAi screens, and QTL mapping to find overlap. These studies included investigations of wild-derived fly populations that identified genes that had correlated expression with metabolic rate, triglyceride storage, trehalose concentration, and pupal weight (Jumbo-Lucioni *et al*. 2010; Williams *et al*. 2015; Scott Chialvo *et al*. 2016), candidate genes from other QTL mapping studies in flies for food intake, weight, mitochondrial phosphate:oxygen (P:O) ratio, and starvation response (Jumbo-Lucioni *et al*. 2010; 2012; Mackay *et al*. 2012; Garlapow *et al*. 2015; Vonesch *et al*. 2016), genes with diet associated eQTLs (Stanley *et al*. 2017; Ng’oma *et al*. 2020), candidate genes with epistatic interactions on triglyceride levels (De Luca *et al*. 2005), and genes showing changes in triglyceride storage when knocked-down by RNAi (Pospisilik *et al*. 2010; Baumbach *et al*. 2014). The complete results from the overlap analyses are given in Supplemental Tables S1-S2.

### Candidate Gene Functional Testing

#### Selection of Genes for Functional Testing

Based on preliminary mapping analyses using models not fully optimized for all confounding and interacting effects, four triglyceride genotype-by-diet QTL with a substantial genetic signal were chosen for further functional testing. We prioritized protein coding genes with high negative log p-values in the mapping results and a functional relationship to lipid metabolism and/or the presence of non-synonymous SNPs within the DSPR Population A (Table S3; McLaren *et al*. 2010; St Pierre *et al*. 2014). We selected a total of 22 genes for gene expression analysis and a subset of 10 genes for mutant characterization (Table S3). The subset for mutant characterization was limited by stock availability and difficulties transferring mutant loci into a common genetic background.

#### Gene Expression Analysis

RNA expression levels were measured using qPCR for the 22 genes selected for expression analysis (Table S3). Thirty-two crosses were selected based on extreme diet effects and 12 crosses were selected for their extreme genetic effect. Fluorescence data was adjusted for reaction efficiencies and normalized to reference genes (Supplemental Methods, File S1). Following a log transformation, an ANOVA was performed to determine significance levels and then corrected for multiple-testing using the Bejamini-Hochberg procedure (Supplemental Methods, File S1; Benjamini and Hochberg 1995).

#### Mutant Characterization

Mutants for 11 genes of the 22 chosen for gene expression analysis were obtained from Bloomington Stock Center based on stock availability (Table S4; Bloomington, IN, USA). For UAS-GAL4-RNAi strains, an Actin5C-GAL4 driver was used to ubiquitously drive transgene expression. Standard breeding methods were used to integrate the transgene from each mutant line into a common genetic background (*W1118*). Up to fifteen samples of ten third instar larvae were collected and tested for triglyceride levels using the same method described above. Triglyceride measurements were adjusted for protein content using the Bradford method (Bradford 1976). Statistical comparisons were performed using standard approaches (Supplemental Methods, File S1).

## RESULTS

We tested Population A of the DSPR RILs (King 2012 a,b) for several phenotypes including male and female pupal weights, and larval triglyceride and trehalose levels in response to a normal laboratory diet and a high fat diet using a round-robin crossing design. The contribution of genetics to phenotype is apparent for all traits (Figure S1, Table S5-S6), with greater variance explained by genotype and genotype-by-diet interactions than by diet alone, as has been observed in previous studies (Reed *et al*. 2010; 2014). We tested six non-null hierarchical mapping models: three for main genetic effects (additive, dominance, founder-by-parent (full)), and each of those effects interacting with diet (additive-by-diet, dominance-by-diet, and full-by-diet). We also mapped overall genetic (main) and genotype-by-diet interaction (diet) effects, which included all the sources of genetic variance (Supplemental Methods, File S1).

### Non-Epistatic Main Effect QTL Analysis

We identified peaks exceeding the global significance threshold (-log p-value > 6) for all tests in male (70 main effect, 17 genotype-by-diet effect) and female (9 main effect, 1 genotype-by-diet effect) pupal weight and for the diet effect on trehalose levels (genotype-by-diet 11; Figures 3, 4, S2-S5, Tables S7-S11). The associations show inflation based on the QQ plots (Figures S6-S9); therefore permutation testing was performed to assess appropriate significance levels for the male weight phenotype (the trait showing the largest number of associations; Table S7, Supplemental Methods, File S1). The overrepresentation of male weight QTL may be due to lower sampling error with larger sample sizes and lower innate within genotype variability in the trait (Reed *et al*. 2010). Given the stringency of negative log p-value >6, and the lack of significant QTLs for some phenotypes, we also considered peaks that were suggestive to allow for some examination of the potential genetic architecture of these phenotypes despite risk of false positives. We found additional suggestive peaks with negative log p-value >2 for triglyceride main (13) and diet (18) effects and for trehalose main effects (19) and investigated these suggestive peaks for the other phenotypes (male weight diet (7), female weight main (11) and diet (18), trehalose main (19) and diet (4); Figures 3, 4, S2-S5, Tables S7-S11). Overall, we characterized 196 unique significant and suggestive non-epistatic peaks. We observed two strong overall patterns in these data: (1) the peaks were non-overlapping across traits (*e.g.* the significant associations for one trait did not predict the significant association in any other trait, Table S7), and (2) there was almost no overlap of the significant peaks between the main and diet effect QTL (Figure 4); the one exception being QTL-index 9396 for male weight (Table S7). There was also one suggestive peak (QTL-index 584) shared between female weight main and diet effect models. The relative variance explained for each model at each QTL and all phenotypes are shown in Figure 5.

**Figure 3:**
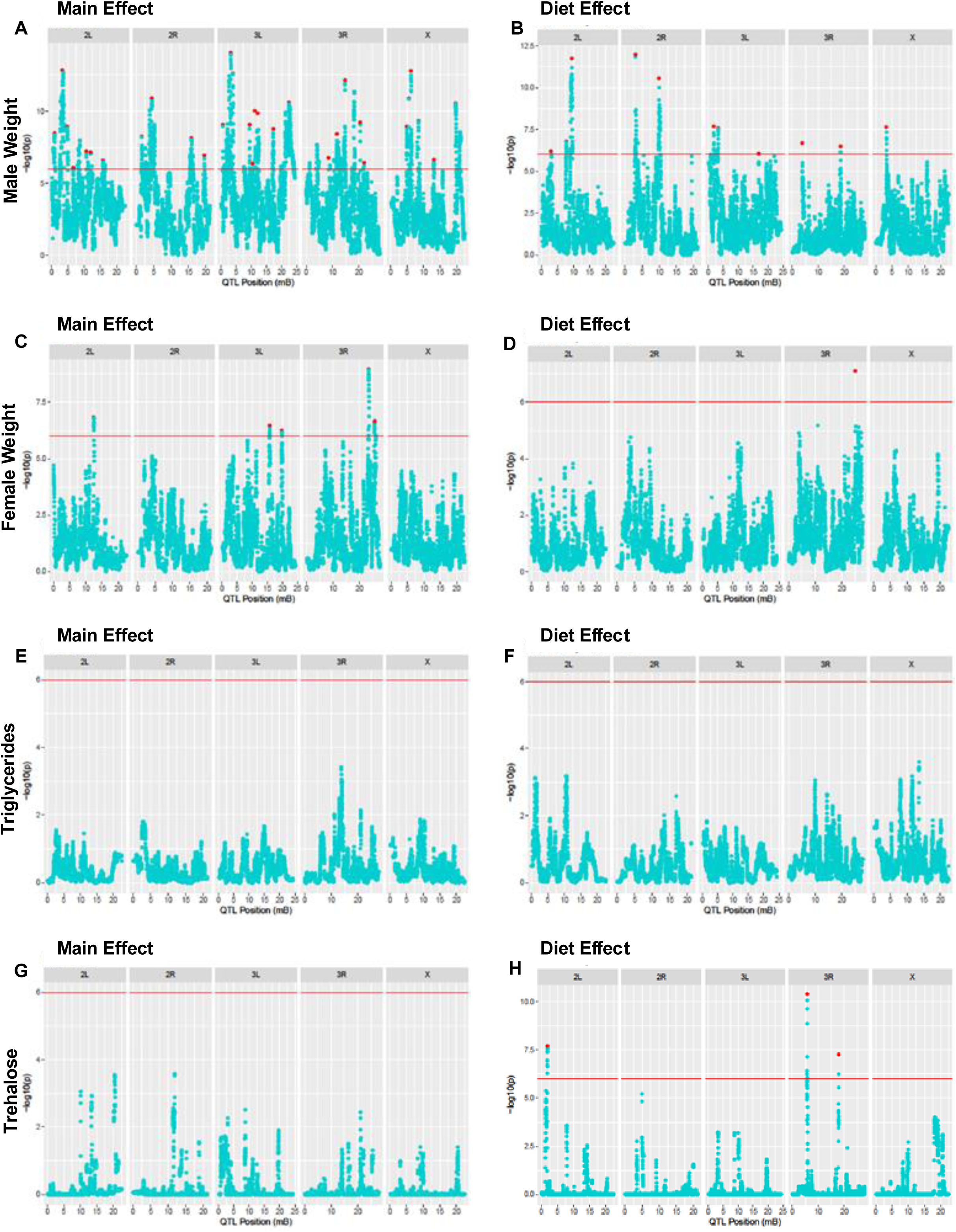
Non-epistatic main and diet effect Manhattan plots for all phenotypes. Manhattan plots showing the negative log p-value for the main effect (full model compared to the null model) for the complete genome in the left-hand panels, while the right-hand panels indicate the diet effect (full-by-diet model compared to the main effect model), for male weight (A, B), female weight (C, D), triglycerides (E, F), and trehalose (G, H). Red lines indicate the significance threshold and local maxima peaks are indicated by red dots. Data available in Tables S7-S11.

**Figure 4:**
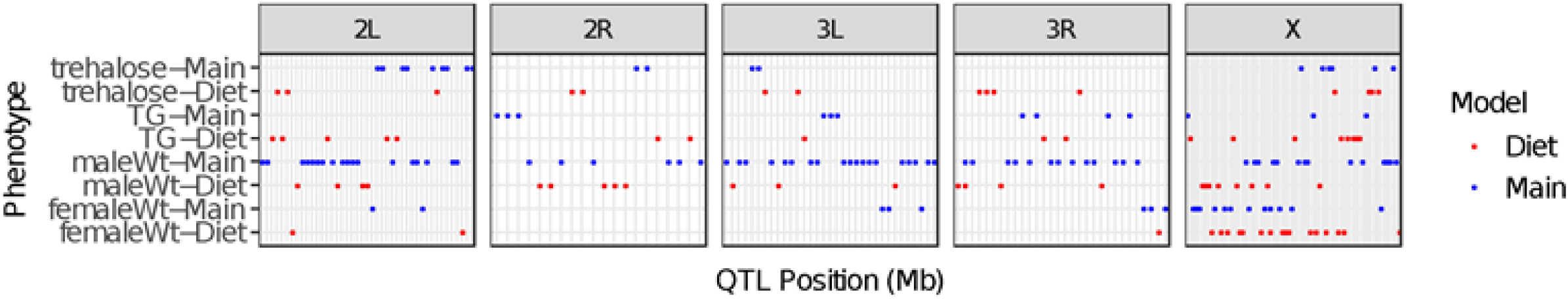
Comparison of non-epistatic peaks (both those defined as significant at negative log p-value (NLP) >6 and suggestive NLP>2) between all phenotypes. For main (blue) and diet (red) models, a point is shown where QTL local peaks are present for each phenotype across the whole genome. These peaks include the significant peaks in Figure 3 but are redrawn here to highlight the lack of overlap of peaks between models and phenotypes. Data in Table S7.

**Figure 5:**
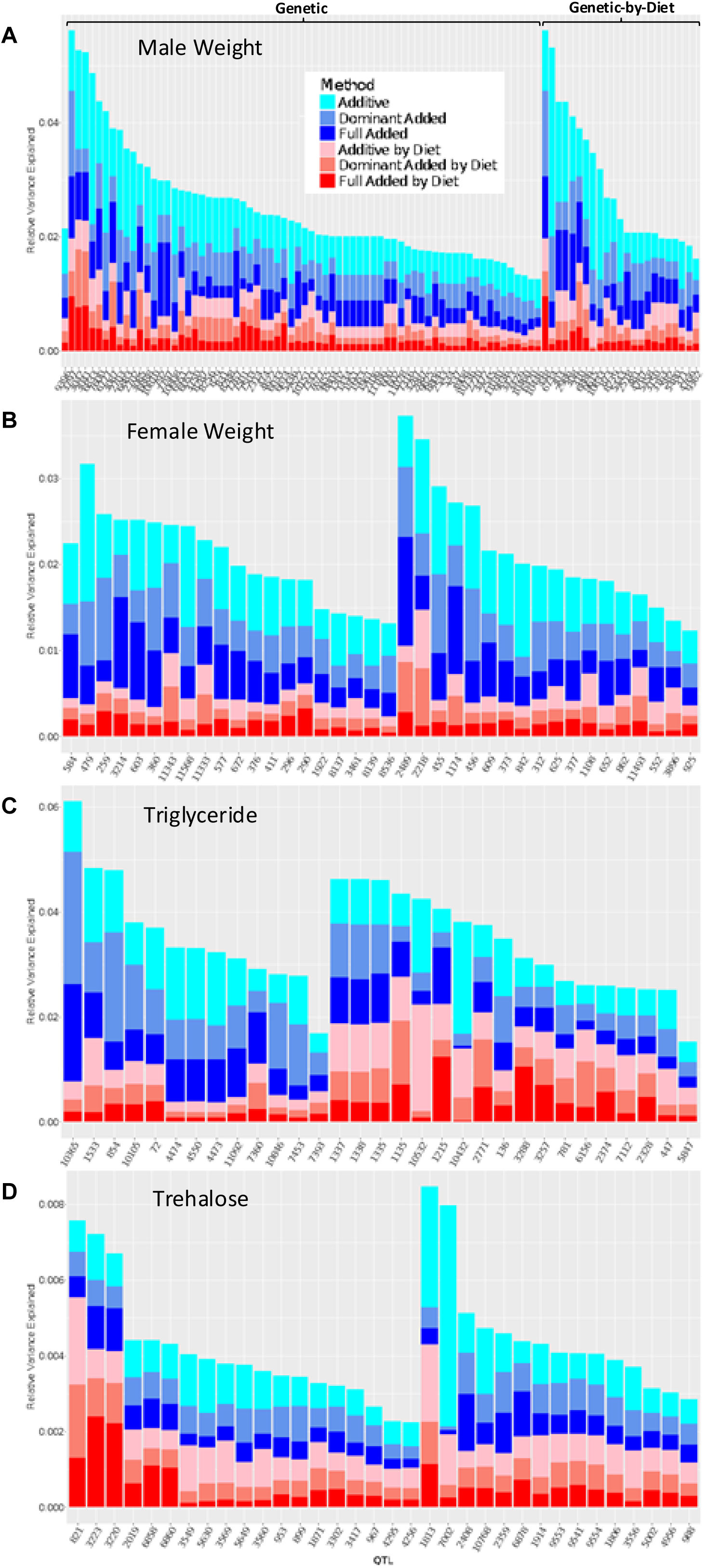
Relative variance explained by each non-epistatic model for all phenotypes. The variance explained by each model for male weight (A), female weight (B), Triglyceride (C), and Trehalose (D). The peaks are ordered based on the cumulative variance explained by all models. The genetic peaks (additive, dominant, full, or main effect models) are displayed toward the left and the genetic-by-diet peaks (additive-by-diet, dominant-by-diet, full-by-diet, or diet model) are displayed toward the right. Peaks identified by the genetic effects models on average explained 1.7%, 1.6%, 2.7%, and 0.2% of variance where the gene-by-diet effects explained 2.1%, 1.6%, 0.9%, and 0.3% of the variance for the male weight, female weight, triglyceride, and trehalose phenotypes, respectively. For the peaks identified by the gene-by-diet interaction model, the interaction effects explain on average 0.8%, 0.6%, 1.6%, and 0.2% of the variance, whereas the main effects explain on average 0.7%, 0.5%, 1.7%, and 0.2% of the variance for male weight, female weight, triglyceride, and trehalose phenotypes, respectively. The raw values of variance explained are listed in Table S5.

### Epistasis QTL Analysis

The epistatic analysis was conducted in an exhaustive manner, thus every pairwise locus combination was tested. Figures S10-S13 show heat maps of the negative log p-value for main epistatic effect (a) and epistatic-diet interaction effect (b), for the male weight, female weight, triglyceride, and trehalose phenotypes, respectively. QQ plots were also generated for the epistatic analyses, and we found that for all but triglyceride main and both trehalose models, the distribution of significance values were consistent with expectation or even deflated (Figure S14). An interaction is considered significant with a negative log p-value greater than 6.0. This threshold results in 8892, 1300, 46, and zero interactions for the main epistatic and 129, 16, 142, and 3386 interactions for the epistatic-diet interaction effects, for male weight, female weight, triglyceride, and trehalose phenotypes, respectively (Table S12, S13). The vast majority of the epistatic loci (98.1%) did not overlap with loci identified in the non-epistatic mapping, and 82.7% of loci were unique to one epistatic model. Only male and female weight had epistatic loci shared across the main and diet effect models (QTL-index 76 and QTL-index 12 loci, respectively), and QTL-index 867 loci was shared across two or more phenotypes (Table S14).

Adjacent epistatic loci in the pairwise-interaction analysis approach were grouped due to their non-independence, which resulted in 68, 63, 16, and zero QTL groups for the main epistatic model, and 28, 7, 32, and 62 QTL groups for the diet epistatic model for the male weight, female weight, triglyceride, and trehalose, respectively (Tables S12, S15). Interaction networks among the groups (nodes), as displayed in Figure 6 and Table S15, showed that some epistatic genomic regions are epistatic “hubs”, which are characterized by many edges connected to that group, while most other groups have few interactions, being epistatic “spokes”. This network structure is typical for biological interaction networks (Barabási and Oltvai 2004; Lu *et al*. 2007). The main epistatic effects for male and female weight, and the diet epistatic effects for the trehalose phenotype were extensive. The large diet epistatic effect for trehalose (Figure 6) was surprising since there were no significant main epistatic effects for the trehalose phenotype. Finally, in contrast to the individually mapped loci, we found that many of the grouped nodes (165 out of 276) were present in multiple phenotypes or models (Table S14).

**Figure 6:**
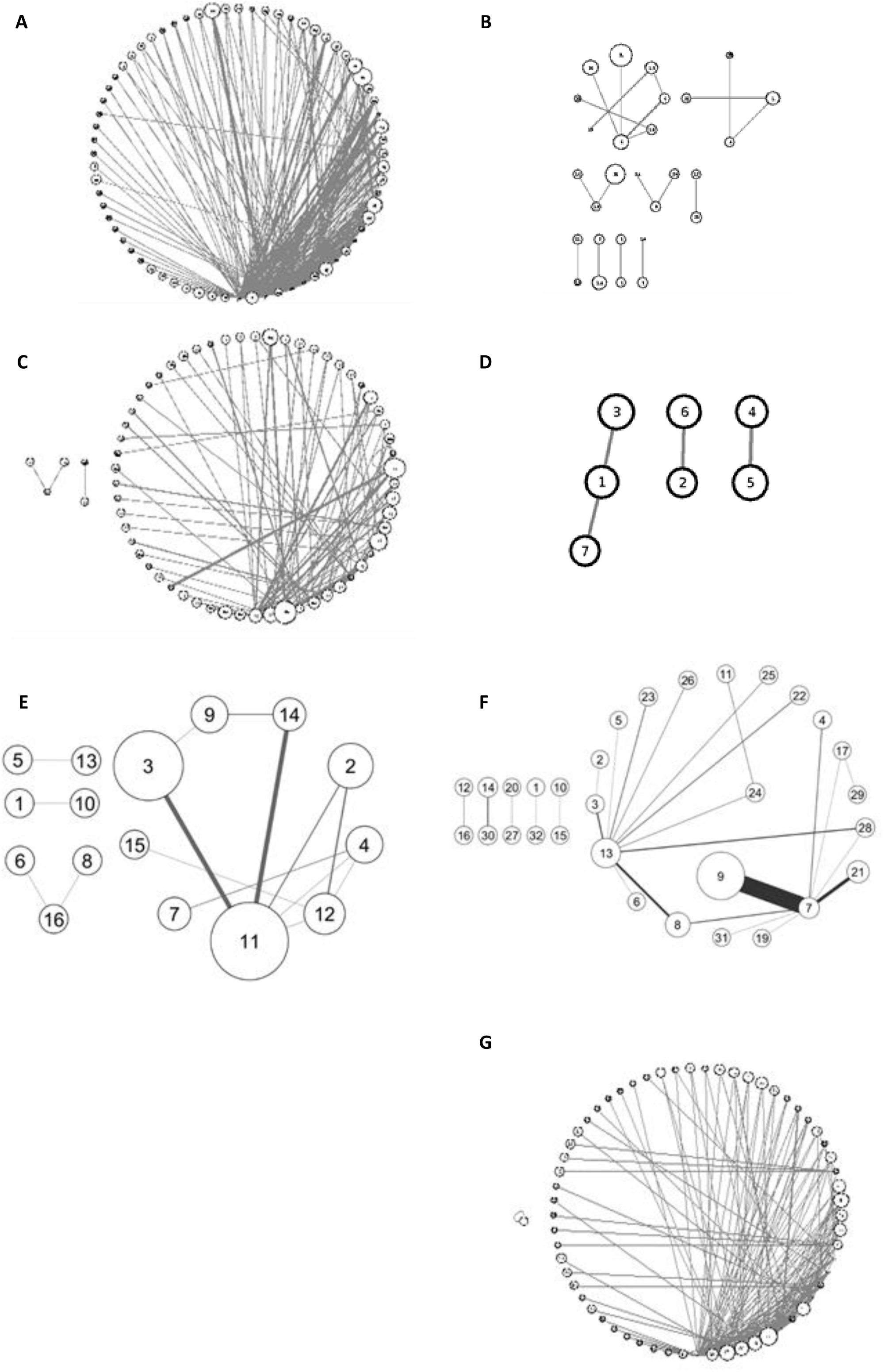
Epistatic interactions for metabolic traits throughout the genome. The QTLs involved in epistatic interactions are grouped based on QTL position. The size of the node depends on the number of QTLs in that group (larger circles indicate more QTLs). The width of the edge depends on the number of significant QTL interactions between QTLs of those groups. Male weight epistatic main effects (A) and diet effects (B). Female weight epistatic main effects (C) and diet effects (D). Triglyceride epistatic main effects (E) and diet effects (F). And trehalose diet effects (G), no significant main effect epistatic interactions for trehalose were observed. The number of significant interactions varied across phenotypes and models suggesting a distinct and complex architecture for each of these traits. All interactions are listed in Tables S14-S15.

### Candidate Genes

From the significant and suggestive QTLs for the non-epistatic effect, we identified 148 unique genes directly overlapping the peaks, an additional 47 closest genes (within 10 kb on either side of the peak), and 8 loci with no annotated gene within 10 kb (Table S16-S17). Thus 28% of the unique loci/peaks (55/196) fell outside of a coding gene span. Interestingly, 14 of the overlapping and closest genes were non-coding RNAs (Table S16-S17). Two genes were shared between male and female weight, *dunce* (*dnc*) and the non-coding RNA *CR44833*. *Proc-R* was shared between male weight and triglyceride storage; and two genes were shared between the main and diet models for male and female weight, *CG9626* and *Nep1,* respectively (Table S16-S17).

For the main epistatic effects, a total of 691, 422, 33, and zero unique candidate genes were identified, while 68, 25, 90, and 700 genes with epistatic dietary effects were found for the phenotypes of male weight, female weight, triglycerides, and trehalose phenotypes, respectively (Table S18). Of all the unique epistatic candidate genes (1655), 84 were non-coding RNAs.

More than half (64.9%) of the significant epistatic peaks were not within the span of protein coding or non-coding RNA genes (2978 out of a total 4582). When the epistatic groups were assessed, 55 of the total 165 groups had no overlap with any genes, and the maximum number of genes assigned to a group was 46 (Table S19). We found over 300 candidate genes that were detected in two or more phenotypes across both the main and diet effect epistatic models (Table S20).

### Gene Ontology Enrichment Analysis

We looked at candidate genes for enrichment patterns relating to biological processes or functions to develop hypotheses of how the detected QTL may be controlling the mapped traits.

#### Non-Epistatic Genes

Gene ontology enrichment analysis of the genes overlapping significant or suggestive non-epistatic peaks across all phenotypes and models (148 genes) showed that biological functions related to post-embryonic development and cell-cell adhesion were overrepresented (Table S21). Within the subset of these genes specific to the genotype-by-diet interactions, development (particularly of the nervous system) was especially enriched (Table S21), while the only distinctive characteristic of the main effect genes was enrichment for the genes having *Snail* transcription factor binding motifs in the regulatory regions. For the phenotype of male weight, there was enrichment for growth/development and rhythmic processes. When we limited our enrichment analyses to only genes found at peaks that survived the permutation testing for male weight (Table S7), we found no qualitative change in the enrichment (*e.g*., regulation of developmental process was significantly enriched in both sets; Table S21), suggesting that the overall enrichment patterns are maintained across varying significance thresholds. Overall, QTLs for female weight were enriched for establishment or maintenance of cell polarity function, as well as cell-cell junction specific expression and genes that interact with microRNA *miR-316*. In addition, we observed a small enrichment for genes that map to the Wnt signaling pathway within the diet specific QTLs for female weight (Table S21). Interestingly, of the 22 genes analyzed for all non-epistatic triglyceride QTLs, two genes were annotated as having the transcription factor binding site for *Adf transcription factor 1* (*Adf-1*) within 2 kb of their transcription start site. The enrichment analysis of the 51 human homologs showed very similar patterns to that seen in the *Drosophila* genes including a number of microRNA interacting gene groups (Table S22).

#### Epistatic Gene Ontology Enrichment Analyses

Across the epistatic genes, there was substantial enrichment for developmental functions, especially in post-embryonic organs and the nervous system, driven by the candidate genes for both male and female weight, and trehalose traits (Table S23, S24), with no obvious qualitative differences in the enriched functions between main and diet epistatic QTLs. A substantial number of the epistatic genes across the phenotypes were enriched for the upstream transcription factor binding motifs of *Adf-1, Mad, GAGA, Snail*, and *Grainy head*. Interestingly, of these transcription factors only the *Grainy head* gene is on the list of epistatic candidate genes. Within the 432 female weight associated epistatic candidate genes, there were 22 with a predicted interaction with *miR-312*.

Many candidate genes for triglyceride epistatic QTLs related to neuron development and included genes in the neuroactive ligand-receptor interaction pathway (LRI, Table S23, S24). Previous research reported that genes involved in LRI might be involved in obesity development (Das and Rao 2007; Almén *et al*. 2014). Two of the LRI associated genes code for 5-hydroxytryptamine (serotonin) receptors, *5-HT1A* and *5-HT1B*, which are involved in feeding behavior (Vickers and Dourish 2004) and their expression was positively correlated with obesity in rats and humans (Voigt *et al*. 2002; Das and Rao 2007). Another LRI gene in this enriched group is *dopamine 2 (D2)-like receptor* (*Dop2R*), and the decrease of D2 receptor levels correlates with increased feeding behavior and obesity (Fetissov *et al*. 2002; THANOS *et al*. 2008; Johnson and Kenny 2010).

Other triglyceride candidate genes of interest include four candidate genes involved in the endocytosis pathway and/or the advanced glycation end products receptor (AGE-RAGE) signaling pathway (which is directly involved in diabetes pathologies such as oxidative stress and diabetic nephropathy; Tan *et al*. 2007; Yan *et al*. 2009). These genes included: *baboon* (*ATR-1*), reported to be involved in glucose metabolism and related to diabetes incidence (Ugrankar *et al*. 2015); *shrub* which negatively regulates the Notch signaling pathway, an important metabolism regulation pathway (Hori *et al*. 2011; Schneider *et al*. 2013; Bi and Kuang 2015); S*WIP* which codes for the polypeptide of the WASH complex (Ropers *et al*. 2011); and *small wing* codes for a phospholipase type C that participates in insulin signaling and has been reported to be involved in metabolic signaling by insulin and the stimulation of glucose uptake (Kayali *et al*. 1998; Lorenzo *et al*. 2002; Murillo-Maldonado *et al*. 2011). For the 337 genes found in two or more models or phenotypes, there was enrichment primarily for organ developmental processes including in the nervous system (Table S25, S26), a reflection of the enrichment pattern among the full list of epistatic candidate genes.

### Overlap of candidate genes with other studies

Ninety-six of the 182 main effect candidate genes overlapped with genes identified in at least one other *Drosophila* study (De Luca *et al*. 2005; Pospisilik *et al*. 2010; Jumbo-Lucioni *et al*. 2012; Mackay *et al*. 2012; Baumbach *et al*. 2014; Reed *et al*. 2014; Garlapow *et al*. 2015; Williams *et al*. 2015; Vonesch *et al*. 2016; Scott Chialvo *et al*. 2016; Jumbo-Lucioni *et al*. 2010a, Ng’oma. 2020) for playing a role in metabolic phenotypes (Table S1). Of these, 34 were found in two or more other studies (Table 1). Of those 34 genes, 10 have been mapped in QTL studies of metabolic phenotypes (Jumbo-Lucioni *et al*. 2012; Mackay *et al*. 2012; Garlapow *et al*. 2015; Jumbo-Lucioni *et al*. 2010a) and 10 have been mapped as epistatic candidate genes for triglyceride storage (Table 1; De Luca *et al*. 2005)). Twenty-nine of these 34 genes showed expression level variation in correlation with one or more metabolic phenotypes (Reed *et al*. 2014; Williams *et al*. 2015; Scott Chialvo *et al*. 2016) and five show a change in triglyceride storage when knocked-down by RNAi in *Drosophila* ( (Pospisilik *et al*. 2010), Table 1). Eighteen were associated with diet-interacting expression QTLs (Ng’oma *et al*, 2020). The complete list of 96 overlapping genes showed enrichment for morphological developmental functions (Table S27). Genes with the transcription factor-binding site for *Trl* (*Trithorax-like*) were enriched in the complete list of overlapping genes (13 genes, Table S27). *Trl* is a transcriptional activator that binds DNA to modify chromatin structure to allow other transcription factors to bind (Chopra *et al*. 2008).

**Table 1:**
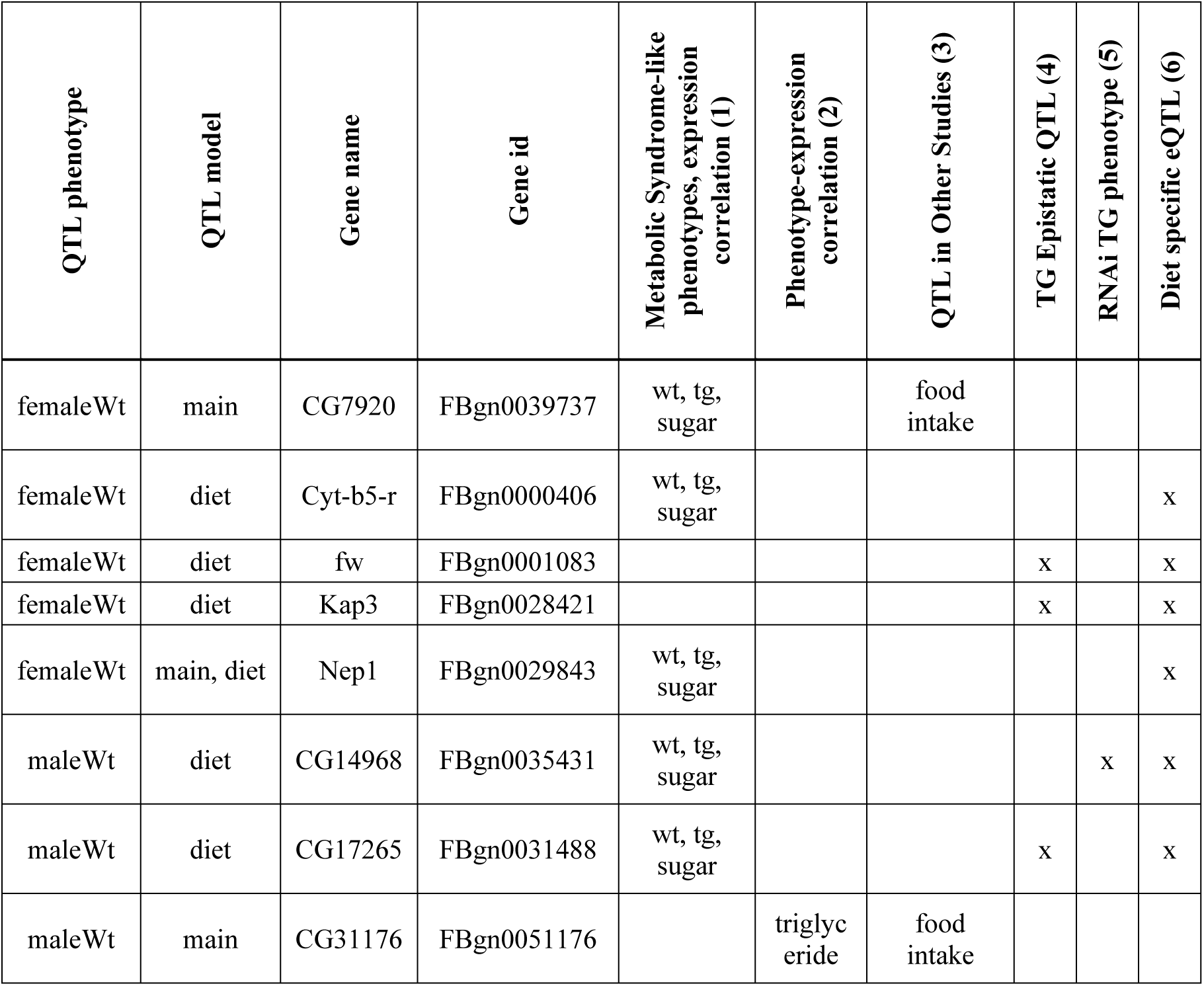
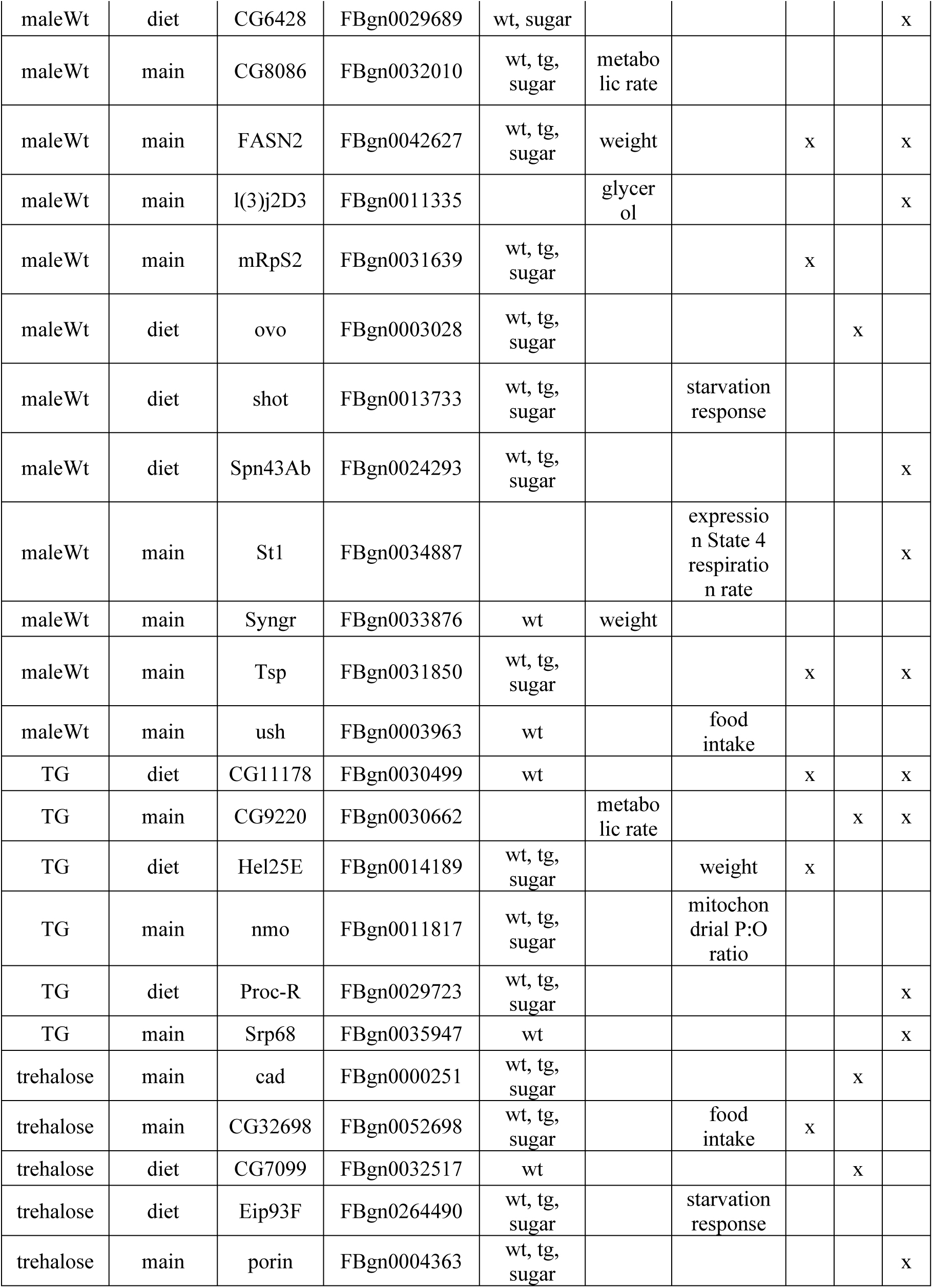
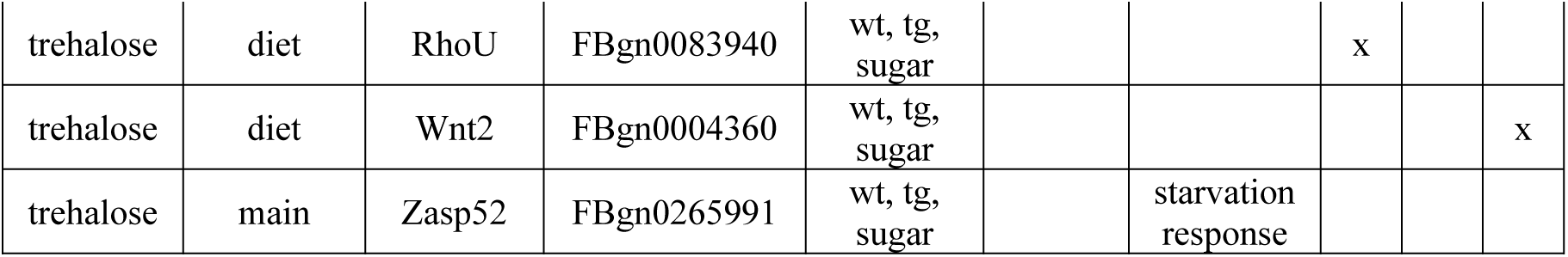
Candidate genes from non-epistatic QTLs with overlap of two or more other high-throughput studies of metabolic traits in *Drosophila*. QTL phenotype- the mapped trait in this study. QTL model - main effect or diet interacting mapping model. Metabolic Syndrome-like phenotypes, expression correlation-phenotypes with gene expression correlation from Reed *et al*. 2014, Williams *et al*. 2015, and Scott Chialvo *et al*. 2016 (1). Phenotype-expression correlation - phenotypes with gene expression correlation from Jumbo *et al*. 2010 (2). QTL in Other Studies - phenotypes mapped to QTLs with the candidate gene from Jumbo *et al*. 2010, Jumbo *et al*. 2012, MacKay *et al*. 2012, Garlapow *et al*. 2015 (3). TG epistatic QTL - candidate genes with QTLs mapped as having epistatic effects on triglyceride storage from DeLuca *et al*. 2005 (4). RNAi TG phenotype - Triglyceride storage phenotype impacted by RNAi knock-down of the candidate gene from Pospisilik *et al*. 2010 (5). Diet specific eQTL - differential expression QTLs associated with diet from Ng’oma *et al*. 2020 (6).

Many of the epistatic candidate genes across all of the phenotypes and models (918) overlapped with genes identified in other screens for metabolic phenotypes (De Luca *et al*. 2005; Pospisilik *et al*. 2010; Jumbo-Lucioni *et al*. 2012; Mackay *et al*. 2012; Baumbach *et al*. 2014; Reed *et al*. 2014; Garlapow *et al*. 2015; Williams *et al*. 2015; Vonesch *et al*. 2016; Scott Chialvo *et al*. 2016; Jumbo-Lucioni *et al*. 2010a; Stanley *et al*. 2017, Ng’oma *et al*. 2020) (Table S2), and again, developmental processes and neuro-muscular functions are some of most enriched biological processes (Table S28). There was also substantial enrichment for genes with GAGA transcription factor binding sites (Table S28). The enrichment patterns remain similar when we consider the 360 epistatic candidate genes in our study that were found to overlap with two or more other studies (Table S29). The 44 epistatic candidate genes that overlapped with candidate genes found in three or more metabolic studies in flies (Table 2), showed significant enrichment for cell migration functions (Table S30). Thirteen of these candidate genes have been mapped as main effect QTLs for metabolic traits in other fly populations (Jumbo-Lucioni *et al*. 2012; Garlapow *et al*. 2015; Jumbo-Lucioni *et al*. 2010) and 17 were previously mapped as having an epistatic effect on triglyceride levels (Table 2, De Luca *et al*. 2005). Forty of these genes have been shown to vary in expression level in correlation with body weight in flies, and many of them also show significant expression variation with triglyceride and trehalose levels (Table 2; Reed *et al*. 2014; Williams *et al*. 2015; Scott Chialvo *et al*. 2016; Jumbo-Lucioni *et al*. 2010). Sixteen of these genes also show a significant change in triglyceride levels with RNAi (Table 2; Pospisilik *et al*. 2010), and 34 of these had diet-specific expression QTLs (Ng’oma *et al*. 2020).

**Table 2:**
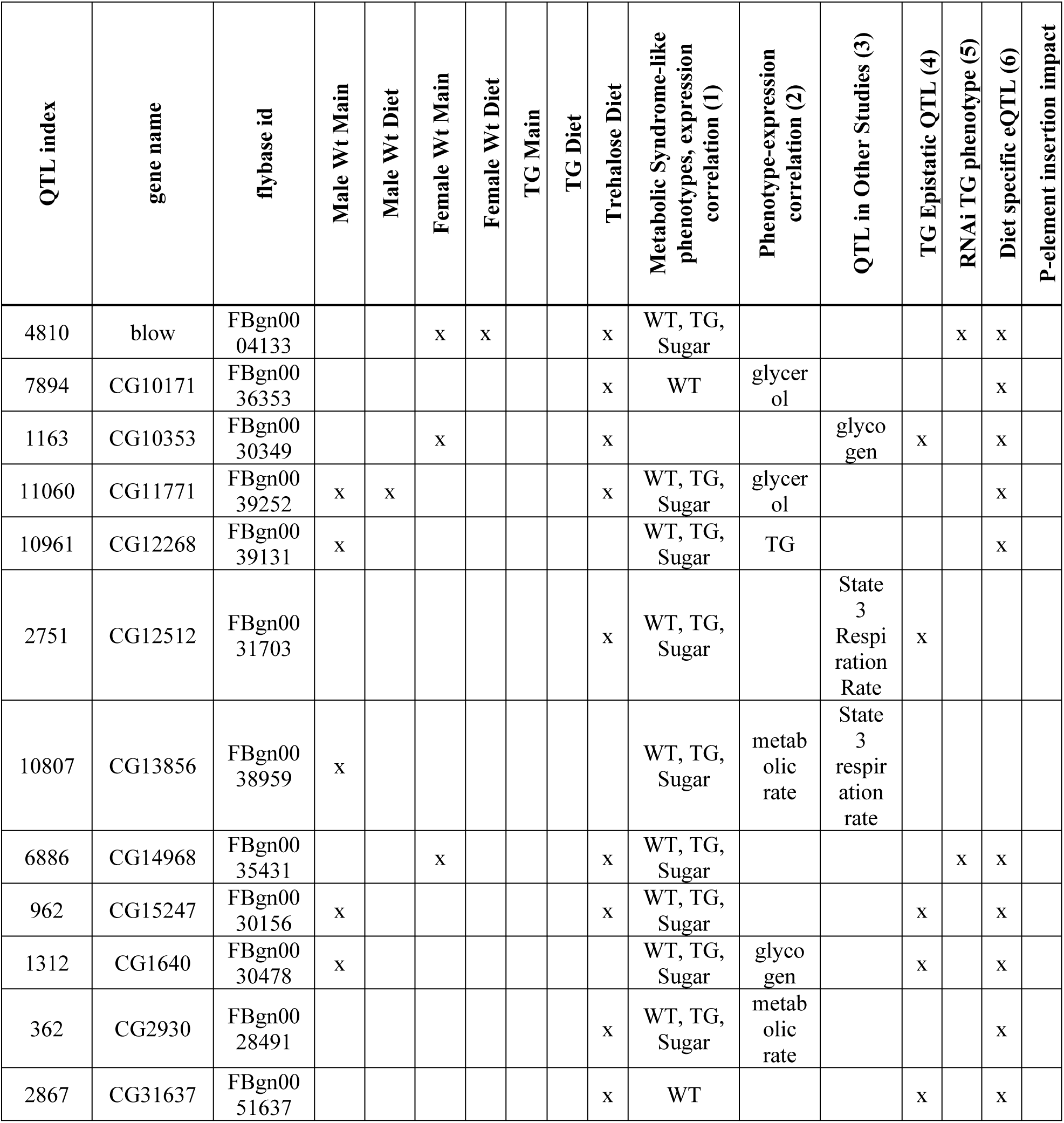
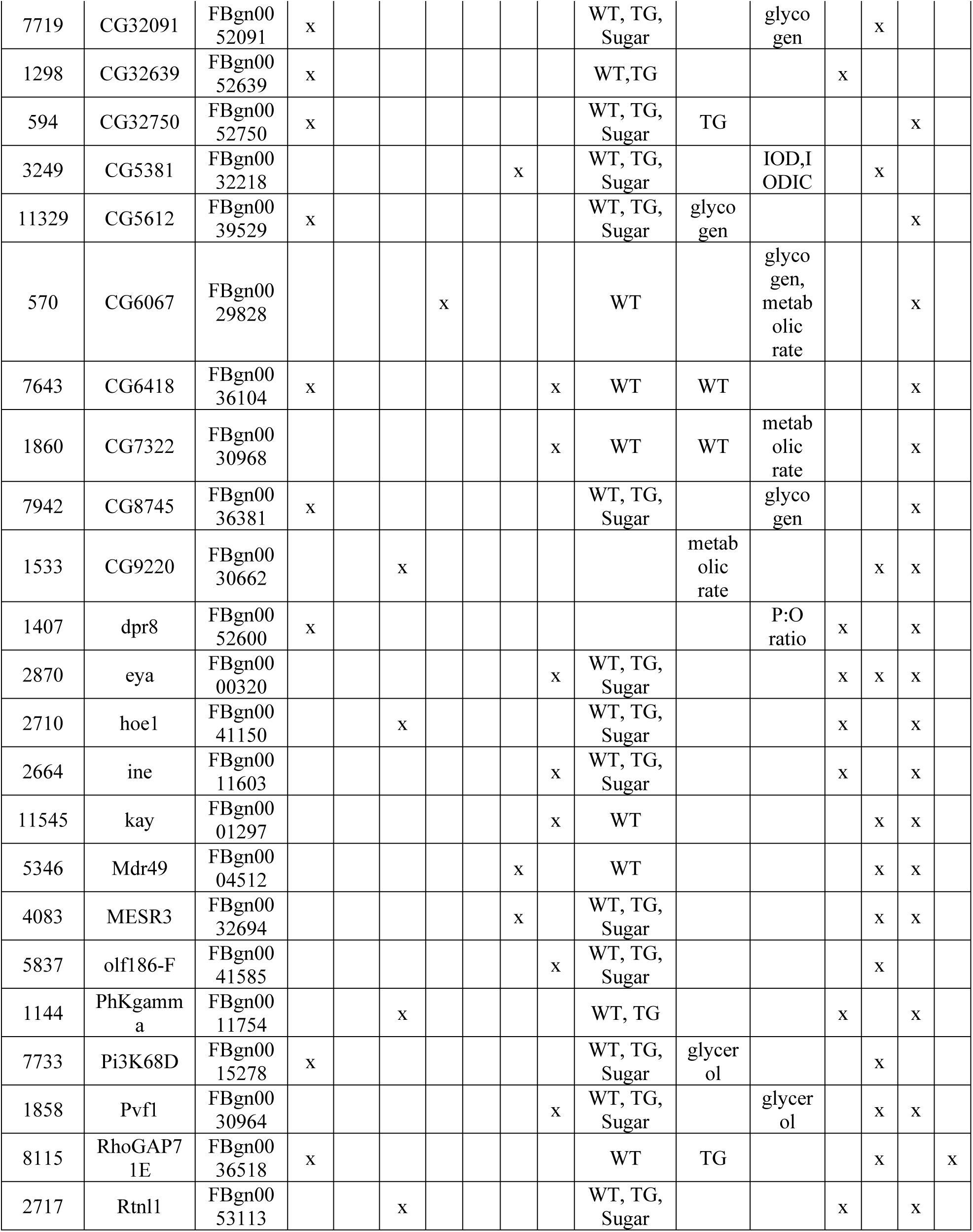
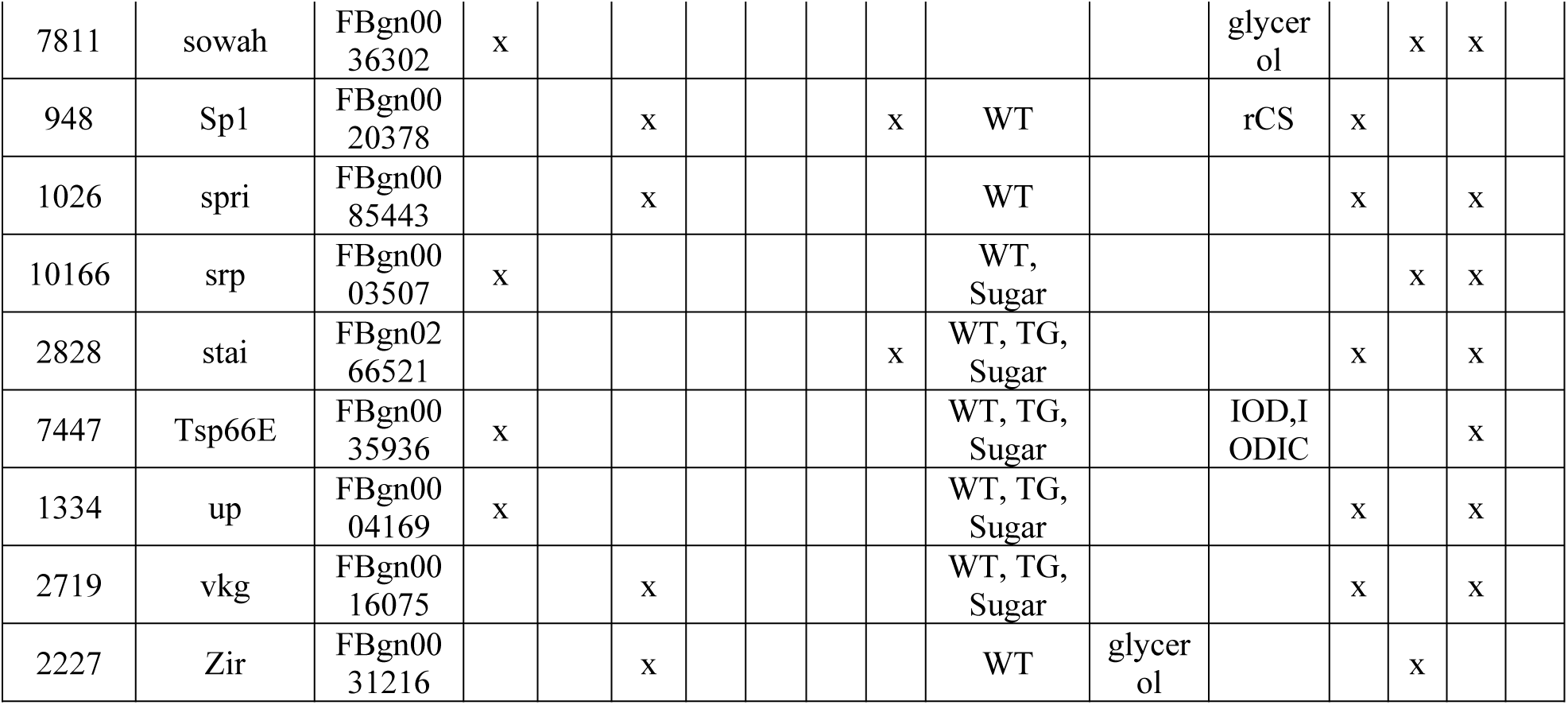
Candidate genes from epistatic QTLs with overlap of three or more other high-throughput studies of metabolic traits in *Drosophila*. QTL index – unique position identified in mapping algorithms, Epistatic QTL phenotypes – association between main or diet-interacting epistatic QTLs and candidate genes for particular phenotypes mapped in this study. Metabolic Syndrome-like phenotypes, expression correlation-phenotypes with gene expression correlation from Reed *et al*. 2014, Williams *et al*. 2015, and Scott Chialvo *et al*. 2016 (1). Phenotype-expression correlation - phenotypes with gene expression correlation from Jumbo *et al*. 2010 (2). QTL in Other Studies - phenotypes mapped to QTLs with the candidate gene from Jumbo *et al*. 2010, Jumbo *et al*. 2012, MacKay *et al*. 2012, Garlapow *et al*. 2015 (3). TG epistatic QTL - candidate genes with QTLs mapped as having epistatic effects on triglyceride storage from DeLuca *et al*. 2005 (4). RNAi TG phenotype - Triglyceride storage phenotype impacted by RNAi knock-down of the candidate gene from Pospisilik *et al*. 2010 (5). Diet specific eQTL - differential expression QTLs associated with diet from Ng’oma *et al*. 2020 (6). P-element insertions with impact on a triglyceride phenotype - Jumbo *et al*. 2010 (7).

### QTL Candidate Gene Testing

Through preliminary statistical analysis using early-stage mapping models prior to optimizing for all confounding and interaction effects (Table S31), we chose a subset of four QTLs for the triglyceride phenotype for functional testing. We examined all genes within the 2-LOD interval around those loci then selected genes for further functional testing based on prior functional information and the presence of non-synonymous SNPs within our dataset. We chose 22 genes as candidates for further evaluation with gene expression analysis and a subset of 10 genes for mutant characterization based on availability mutant stocks (Tables 3, S32-S34). An FDR of 0.3 was used to determine significance in these analyses. Two candidate genes (*Spt20* and *Liprin-β*) in the dominant diet-interacting QTL on 3L (3L-135) showed functional evidence of having an effect on triglyceride levels. Expression of *Spt20* had a triglyceride-by-diet interaction and the *Liprin-β* knock-out mutant had higher triglyceride levels overall than the wildtype control, but no correlation with triglycerides was observed in gene expression analysis for *Liprin-β* (Figure 7, S15, Tables 3). Two additional genes (*CG2533* and *rho-4*) in a second dominant diet-interacting QTL (X-115) were examined. A double loss-of-function mutation in *CG2533* and *rho-4* had lower triglycerides overall and up-regulation of *rho-4* resulted in a genotype-by-diet interaction driven by the mutant’s reaction to a high fat diet (Figure 7, S15, Tables 3). For the candidate genes tested in the two QTLs (2L-107, 2R-170) identified in the full diet interaction model, *Dpy30L1* expression had a triglyceride-by-diet interaction, the *EMC3* knock-out mutant had elevated triglyceride levels, *Treh* expression had a significant triglyceride-by-diet interaction, and a mutant with upregulated *CG4266* had increased triglyceride levels (Figure 7, S15, Tables 3). In addition, a *Magi* knock-out had higher triglycerides than the control, but the mutant with upregulated *Magi* showed no change (Figure 7, S15, Tables 3). Several controls were also used to ensure the validity of the mutant characterization experiment. A loss-of-function mutation in *pudgy*, a gene previously shown to have an inverse relationship between expression and triglycerides, had decreased triglycerides compared to the control as we would expect (Table S34; Xu *et al*. 2012).

**Table 3:** Significant Functional Triglyceride Candidate Genes. Triglyceride-associated candidate genes for effect on triglyceride storage phenotypes were tested for whether they exhibited changes in expression across diets or showed a change in triglyceride phenotype when mutated. ^a^ FDR<0.3 corrected p-values

| QTL | Gene | qPCR<sup>a</sup> | Mutant Characterization<sup>a</sup> |  |
| --- | --- | --- | --- | --- |
|  |  | Trigly * Diet | Mutation | Mutation * Diet |
| X-115 | rho-4 | ns | 0.0146 | 0.0623 |
|  | CG2533 & rho-4 | - | <.0001 | ns |
| 2L-107 | EMC3 | ns | <.0001 | ns |
|  | Dpy-30L1 | **0.2493** | ns | ns |
| 2R-170 | CG4266 | ns | <.0001 | ns |
|  | Treh | **0.0886** | - | - |
|  | Magi (Knock-out) | - | 0.0006 | ns |
| 3L-135 | Liprin-b | ns | 0.0249 | ns |
|  | Spt20 | **0.0886** | - | - |

**Figure 7:**
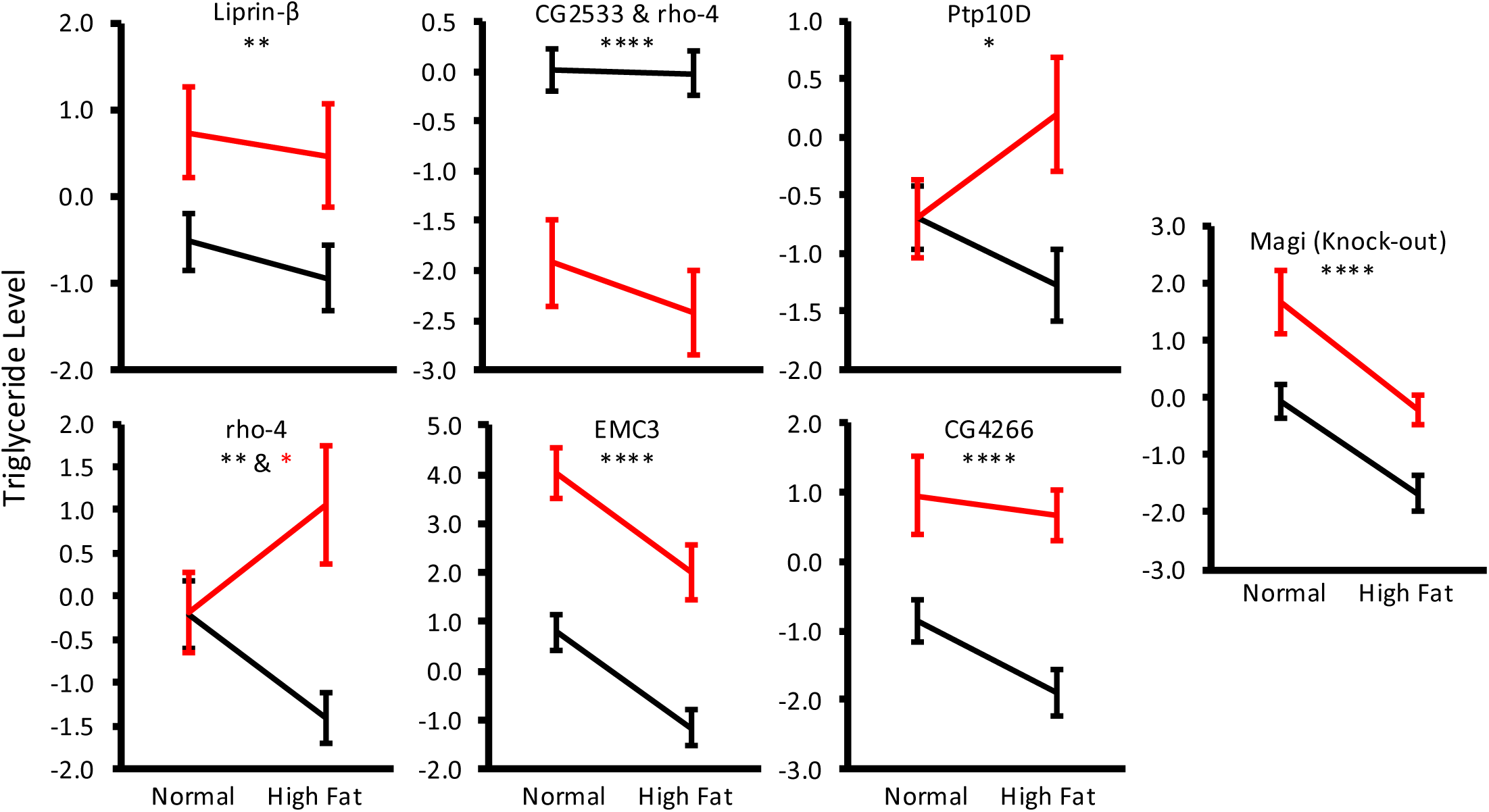
Candidate gene mutant triglycerides. Visualization of relationship between mutant (red) and control (black) triglycerides. All genes shown were significant for genotype (black stars) and one had a genotype-by-diet interaction (red star). FDR<0.3 used as multiple testing correction. 0.1 - 0.05 *, 0.05 – 0.01 **, 0.01 – 0.001 ***, <0.001 **** Data in Table S32.

## DISCUSSION

### Genetic Architecture and Function of Non-Epistatic Loci

One of the most compelling and fundamental findings from overall patterns in this study is there was virtually no overlap among the non-epistatic loci between the main versus genotype-by-diet effects. This is counter to our initial expectation that the genes responsible for population-wide main genetic effects would likely be the ones responsible for the environment-specific regulation of the phenotypes. This suggests that studies that limit mapping analysis to a population in a single environment may be missing much of the genetic variation for a given phenotype in the population, which is supported by the substantial portion of diet specific variance that could be assigned to the main effect loci (Figure 5). This idea could be further tested in the future by studies of this dataset that compare diet-specific main effect mapping to the interacting loci found in this study. Also, this confirms that there is likely substantial cryptic genetic variation that can be exposed if a population transitions to a new environment, providing fresh additive genetic variance on which selection can act (Gibson and Reed 2008). In addition, when we compare the functional enrichment for the main effect versus diet effect loci we do not find common functions between groups suggesting that the types of genes responsible for average versus plastic genetic variation are likely functionally different (Table S21, S23). However, it should be noted that the relatively short lists of candidate genes found in the main effect mapping may limit our ability to make this comparison and further study of these types of comparisons is needed.

One potential oddity to these findings is that there were no significant genotype-by-diet effect QTL when there were significant genotype-by-diet effect QTL for trehalose (13), when the dietary manipulation was based on changing fat content rather than carbohydrate. Given that there were also no significant main effect QTL for triglycerides, one possibility is that the triglyceride phenotype, relative to the trehalose phenotype, harbors relatively little non-epistatic genetic variation (lower heritability). This is consistent with what was observed a different wild-derived fly population tested across four different diets (including a high fat and a high sugar diet) where they found that genetic effects accounted for 10.9% and genotype-by-diet effects accounted for 11.5% of the total phenotypic variation for triglyceride levels (Reed *et al*. 2010). In contrast, for trehalose there was much more phenotypic variance attributed to genetic (23.1%) and genotype-by-diet (17.3%) effects (Reed *et al*. 2010).

In the functional enrichment analysis of the non-epistatic effect candidate genes, we found a number with links to metabolism and metabolic syndrome. For example, the cluster for rhythmic process is intriguing since the circadian clock system is tightly connected with metabolism at molecular and behavioral levels, and alteration of sleep patterns is associated with metabolic disruptions that may lead to adverse adipose accumulation and obesity in humans (Bray and Young 2007; Laposky *et al*. 2008). Among the genes with rhythmic process annotations, FBgn0016694 (*PDP1*) draws special attention since isomers of *Pdp1* are expressed in the nervous system (Pdp1ε) and fat body (Pdp1γ), and loss of expression of *Pdp1* resulted in significantly reduced lipid levels in *Drosophila* (Dzitoyeva and Manev 2013). In addition, five genes of the 50 male main effect QTLs have an upstream binding site for the transcription factor *Snail,* which is involved in tissue differentiation and neuronal development functions (Nieto 2002; Aybar *et al*. 2003; Boutet *et al*. 2006; Tang *et al*. 2016).

A distinctive gene from the non-epistatic gene list, *FASN2* (*fatty acid synthase 2*), is reported to be activated by the presence of dietary sugar and involved in protection against sugar toxicity (Garrido *et al*. 2015). *FASN*, the human ortholog of *FASN2*, encodes a multifunctional enzyme involved in long-chain fatty acid extension and cholesterol biosynthesis, and bears allelic variations in humans associated with obesity (Moreno-Navarrete *et al*. 2009; Schleinitz *et al*. 2010).

Another intriguing set of candidate genes for male weight were enriched for growth function and contained *slpr* (*slipper*) and *Hipk* (*Homeodomain interacting protein kinase*), which are involved in the JNK cascade and Notch positive regulation, respectively. The JNK pathway can disrupt the insulin signaling pathway and thus plays a critical role in the development of obesity and type 2 diabetes (Hirosumi *et al*. 2002). In addition, *Notch* mutants are more resistant to a high fat diet, have better insulin sensitivity, glucose tolerance, and increased energy expenditure (Bi *et al*. 2014).

Two additional genes of interest among the non-epistatic candidate genes include *Dunce* (orthologous to *phosphodiesterase 4D* (*PDE4D*)) and *Octopamine β2 receptor* (orthologous to *beta-2-adrenergic receptor* (*ADRB2*)), which are involved in adenylate cyclase-activating adrenergic receptor signaling pathway (Table S22). Variation in several adrenergic receptors is correlated with weight gain and obesity (Kadowaki *et al*. 1995; Clément *et al*. 1995) and activation of adrenergic receptors in adipose tissue leads to lipolysis in humans (Langin 2006). Human allelic variations of *ADRB2* are also related to blunted accumulation in free fatty acids and glycerol, as well as decreased fat oxidation, independent of obesity, physical activity, and type 2 diabetes (Jocken *et al*. 2007; Kilpeläinen *et al*. 2008; Gjesing *et al*. 2009).

### Main Genetic Effect Candidate Gene Functional Testing

We were able to test candidate genes from four QTL identified in the preliminary analyses of the triglyceride data either through a genotype or a genotype-by-diet interaction. Several of these genes can be mechanistically connected to triglyceride regulation or obesity, while the mechanism for the connection of others is less obvious. For those that have independent associations with obesity or triglyceride regulation, these genes validate our method of locating QTL and determining potential candidate genes. Fourteen of our functionally tested candidates have been identified by other high throughput metabolic studies in flies (Table S35) (De Luca *et al*. 2005; Pospisilik *et al*. 2010; Jumbo-Lucioni *et al*. 2012; Mackay *et al*. 2012; Baumbach *et al*. 2014; Reed *et al*. 2014; Garlapow *et al*. 2015; Williams *et al*. 2015; Vonesch *et al*. 2016; Scott Chialvo *et al*. 2016; Jumbo-Lucioni *et al*. 2010). For example, higher triglycerides in an *EMC3*-knockdown mutant confirms results from a previous study which found *EMC3* to be an ‘anti-obesity’ gene in an RNAi screen that measured fat storage in adult flies (Baumbach *et al*. 2014). Another gene tested in this study, *Trehalase* (*Treh*), encodes the enzyme that breaks down trehalose into two glucose molecules. Trehalase gene expression had a significant triglyceride-by-diet interaction, consistent with the linked regulation of sugar and fat metabolism. The expression level of *Treh* has also been identified as having expression levels correlated with sugar and triglyceride content, as well as body weight across multiple diets in other studies (Reed *et al*. 2014; Williams *et al*. 2015). A member of the Rhomboid family localized to the plasma membrane, *rho-4* (which encodes an intramembrane protease that cleaves and activates *epidermal growth factor receptor* (*EGFR*) ligands (Urban *et al*. 2002), was found to have an effect on triglycerides in two of our mutant tests. It has also been identified as a candidate gene for epistatic effects on triglycerides in another study (De Luca *et al*. 2005).

Expression of *Magi,* a membrane associated with guanylate kinase activity (Schoenherr *et al*. 2012), has been associated with body weight (Reed *et al*. 2014; Williams *et al*. 2015) and is a candidate gene in a QTL for interocular distance - a growth measure (Vonesch *et al*. 2016). We also found that the endogenous expression level of *Dpy-30L1* is positively correlated with triglyceride storage. *Dpy-30L1* is an expendable portion of the H3K4 methylation complex, and was identified as a moderately validated obesity gene by RNAi (Baumbach *et al*. 2014).

We tested multiple candidate genes for each of our functionally tested QTLs (Table S32) and found evidence of significant effects on triglyceride storage for multiple genes for each locus, which supports two, non-exclusive hypotheses. First, the QTLs we observe may be the result of genetic variants in multiple linked genes, and/or traits like triglyceride storage are so polygenic that there is a reasonable chance that if one modifies any given gene experimentally there will be an observed effect on the phenotype. Direct allele swapping (*e.g.* CRISPR-Cas9) for the variants around these genes may be the only direct way to determine with certainty which variants, and associated gene or genes, are responsible for a given QTL. Looking across all four mapped traits, comparisons of candidate genes found in this study to other high--throughput studies of metabolic traits in *Drosophila* (Tables 2, S1 & S16; De Luca *et al*. 2005; Pospisilik *et al*. 2010; Jumbo-Lucioni *et al*. 2012; Mackay *et al*. 2012; Baumbach *et al*. 2014; Reed *et al*. 2014; Garlapow *et al*. 2015; Williams *et al*. 2015; Vonesch *et al*. 2016; Scott Chialvo *et al*. 2016; Jumbo-Lucioni *et al*. 2010, Ng’oma *et al*. 2020), supported a number of additional candidate genes as likely having functional relevance to these metabolic traits.

### Genetic Architecture and Function of Epistatic Loci

As we observed the non-epistatic mapping results, the loci responsible for main epistatic effects were largely non-overlapping with loci showing a diet effect epistatic interaction. In addition, most of the epistatic loci had no non-epistatic effects. This demonstrates, as shown in other studies (Swarup *et al*. 2012; Huang *et al*. 2012; Chen *et al*. 2017), that limiting the analysis of epistatic effects in mapping studies to loci initially detected as having non-epistatic effects will miss the majority of the epistatic loci. The patterns found in the epistatic analyses further emphasize the substantial pool of cryptic genetic variation that is exposed by an environmental change. This important role of environment is most starkly emphasized by the trehalose phenotype that has the complete lack of significant epistatic main effect QTLs but a substantial number of diet effect epistatic QTLs (Figure 6). While statistical significance is not itself “significant” (Gelman and Stern 2006), when comparing across analyses with analysis-specific underlying assumptions, an arbitrary threshold of some type is required to facilitate comparisons. Ideally, future studies will also be able to compare relative effect sizes for epistatic versus non-epistatic loci across in a similar dataset.

The architecture of the network of interactions among groups of epistatic loci was also of interest. We found the overall architecture to be consistent with the hub-and-spoke structure observed in other biological systems (Jeong *et al*. 2000) characterized by a few nodes with many interactions (hubs) and many nodes with few interactions (spokes). It should be noted however that there was a very strong positive correlation (p<0.0001) between the size of the epistatic group and the number of group-to-group pairwise interactions that group participated in (Table S36). Also, the larger groups overlapped with more candidate genes. The question then is whether an increased “size” of an epistatic group is driven by the number of loci that happen to be in proximity that are independently interacting with other loci in the genome, or by a few highly epistatic loci, which would drive the appearance of additional statistical interactions at adjacent loci due to linkage disequilibrium.

In terms of the patterns in gene function, the epistatic QTL showed similar patterns of enrichment across both male and female weight and the trehalose phenotype, with strong enrichment for genes involved in signaling and organismal growth, including transcription factors, transmembrane transporters and receptors, and structural protein binding. Given this pattern of enrichment was observed across three phenotypes and within main effect and diet effect loci, it suggests that genes involved in signaling processes and development might be especially likely to exhibit epistatic variation in populations in general. Patterns in other studies of epistatic loci have shown that “essential” genes, those showing more drastic phenotypic effects when knocked down, are more likely to be involved in epistatic interactions (Phillips 2008), and that epistatic loci also show some enrichment for signaling and development (Wei *et al*. 2012; Huang *et al*. 2012).

Wnt signaling was enriched in the genes associated with main effects (Tables S21-22) and epistatic effects (Table S24). Wnt signaling regulates adipogenesis and its disruption results in transdifferentiation of myoblasts in adipocytes (Ross *et al*. 2000). One Wnt pathway gene found in these lists, *Myc*, is a transcription factor that is expressed in muscles and the fat body and is involved in regulation of glucose metabolism (Ugrankar *et al*. 2015). In addition, across all of the non-epistatic and epistatic effect candidate genes, 88 were in common to both types of models (Table S37). These genes showed enrichment for growth and development functions (including of the nervous system) suggesting that although overall enrichment patterns across different models and genetic architecture differed, there may still be a core set of genes that play a role in MetS-like traits regardless of the explicit architecture of the genetic variation (Table S38). This pattern is consistent with the omnigenetic hypothesis which postulates that a core set of genes will mediate phenotype-specific variation mapped to tissue and development stage specific genes (Boyle *et al*. 2017).

### Regulation and Non-coding QTLs

For both the main effect and the epistatic effect loci, a substantial portion (main 43% and epistatic 42%) mapped to intergenic regions. This highlights the importance of genetic variation in cis-regulatory and enhancer elements in shaping the architecture of complex traits. In addition, there was substantial gene ontology enrichment for transcription factor activity among the protein coding genes associated with the epistatic QTLs, and many of the other candidate genes for both the main and epistatic effect loci were non-coding RNAs.

Another intriguing gene group, from the non-epistatic genes, clustered based on interactions with microRNA *miR-316*, and within this group, *Cyt-b5-r* attracts special attention as it is involved in lipid metabolic processes. The upregulation of *Cyt-b5-r* resulted in significant increase in larval lipids (Zhao *et al*. 2012). Interestingly, glycosylation, unsaturation, and hydroxylation of sphingo lipids during *Drosophila* development is also correlated with *Cyt-b5-r* expression (Guan *et al*. 2013). Further, microRNAs have been shown to affect triglyceride levels (Xu *et al*. 2003; Teleman *et al*. 2006), and to effect within and between species morphological variation in *Drosophila* trichrome patterns (Arif *et al*. 2013). They are also involved in regulating neural development (Tsurudome *et al*. 2010; Yatsenko *et al*. 2014). *miR-316* has a significant effect on lifespan in male flies (Chen *et al*. 2014) and also shows plasticity in expression level in response to age and mating status (Zhou *et al*. 2014). This is especially interesting as these *miR-316* interacting candidate genes also show plastic responses to diet. In addition, three of the 15 genes associated with the diet effect on trehalose levels are predicted to interact with *miR-281* (Table S21). *miR-281* shows plasticity in expression due to age and mating status similar to that seen in *miR-316* (Zhou *et al*. 2014).

Of special note are the 22 *miR-312* interacting genes found for the epistatic female weight loci. These 22 genes (Table S39) are also predicted to interact with other members of the *miR-310* cluster (miR-310, miR-311, and miR-313) as well as *miR-92a* and *mir-92b*, though no QTLs map close to any of these miRNAs. *miR-92a* and *miR-92b* are the ancient paralogs that gave rise recently (within the *Drosophila* genus) to the *miR-310* cluster through duplication, and subsequently, the cluster shows evidence of having experienced a selective sweep (Lu *et al*. 2008). The *miR-310* cluster specifically has been attributed to diet-specific regulation of nutrient storage through the Hedgehog signaling pathway (Çiçek *et al*. 2016). Additionally, it influences nicotine and ethanol sensitivity (Sanchez-Díaz *et al*. 2015; Ghezzi *et al*. 2016), and innate immune responses (Li *et al*. 2017). The strong enrichment for genes likely regulated by *miR-310* cluster microRNAs and non-coding RNA candidate genes draws attention to the important role non-coding genes play in metabolic process. Also, the substantial number of intergenic loci, as well as the transcription factors and signaling function genes among the coding candidate genes, highlights the complex regulatory architecture of phenotypic variation.

A substantial number of loci leading to long lists of candidate genes found in this study leads one to question what general properties of genetic variation for complex traits can be identified. One strategy would be to focus especially on candidate genes that have been identified in association with similar traits across multiple studies. When we did that for the non-epistatic effect loci we found 96 genes that have also been identified as metabolism-trait related in at least one other high-throughput study (Table S1) and found 34 across two or more additional fly studies (Table 1). These shared genes show the same trends in functional enrichment observed across the non-epistatic loci in general, thus giving us no special insight to a most “important” group of genes for metabolic trait variation. Similarly, the epistatic candidate genes highlighted by one or more other studies reflected the same patterns of functional enrichment for morphological and nervous system development observed in the full epistatic candidate gene list. Despite the lack of clarity given by the functional enrichment analysis of these repeatedly implicated genes, the genes themselves are likely still worth more careful examination on an individual level and appear to be important across a variety of metabolic traits. A few of these genes, just by the preponderance of prior evidence of involvement in metabolic processes (*e.g*. *FASN2*), suggest that looking at the intersection among studies will find genes fundamental to variation in metabolic phenotypes.

A comparison of the genes found across multiple studies established another pattern that the association between a gene and particular genetic mapping model or phenotype may only be broadly reflected in other studies. For example, the gene *CG32698* was mapped to be a candidate gene in the main diet trehalose model in this study but was mapped to have an epistatic effect association with triglycerides in another study (DeLuca *et al*. 2005). It was also identified as a main effect candidate gene for food intake (Garlapow *et al*. 2015) and have expression levels correlated with weight, triglycerides, and trehalose (Reed *et al*. 2014, Chialgo *et al*. 2016) in two other independent studies (Table 1). *CG32698* also has no annotated biological function.

Therefore, if we limited the search to only main effect QTLs for trehalose levels, or only focused on loci with particular functional annotations, we would not have seen the broader pattern of *CG32698*, an uncharacterized gene, likely being truly biologically important across metabolic phenotypes.

Returning to our original framing questions, overall, our study shows that the genetic bases of metabolic traits are highly complex and vary substantially between main and dietary-plastic effects. In addition, the loci responsible for direct genetic effects are largely not the ones involved in epistatic interactions for the traits. Furthermore, while there is some commonality of significant loci across studies, most QTLs appear to be unique to a particular population, supporting the omnigenetic hypothesis that there are many genetic paths to arrive at a common phenotype probably mediated through a smaller number of critical hubs (Boyle *et al*. 2017). The general patterns of functional enrichment in the epistatic loci show that genetic variation in development, growth, and neurological functions play a consistent role in influencing phenotypic variation in metabolic traits. As most of the candidate genes identified in this study would not be highly ranked as potentially good candidates *a priori*, our findings add to mounting evidence that a gene-centric view of complex disease is unlikely to produce fundamental understanding of the underlying causes for these conditions (Rockman 2012).

## Data Availability

Raw data generated by this study and the supplemental tables are available through *figshare* (https://doi.org/10.6084/m9.figshare.29817206, https://doi.org/10.6084/m9.figshare.29817683). Further code data is available via GitHub (https://github.com/rmathur87/DLinkMaP/tree/master).

## ACKNOWLEDGEMENTS

This work was made possible in part by a grant of high-performance computing resources and technical support from the Alabama Supercomputer Authority.

## FUNDING

Support for this study was provided by R01GM098856 (to LKR and AMR) and R25GM130517 (to LKR). This work was made possible in part by a grant of high-performance computing resources and technical support from the Alabama Supercomputer Authority.

---

**Figure S1:**
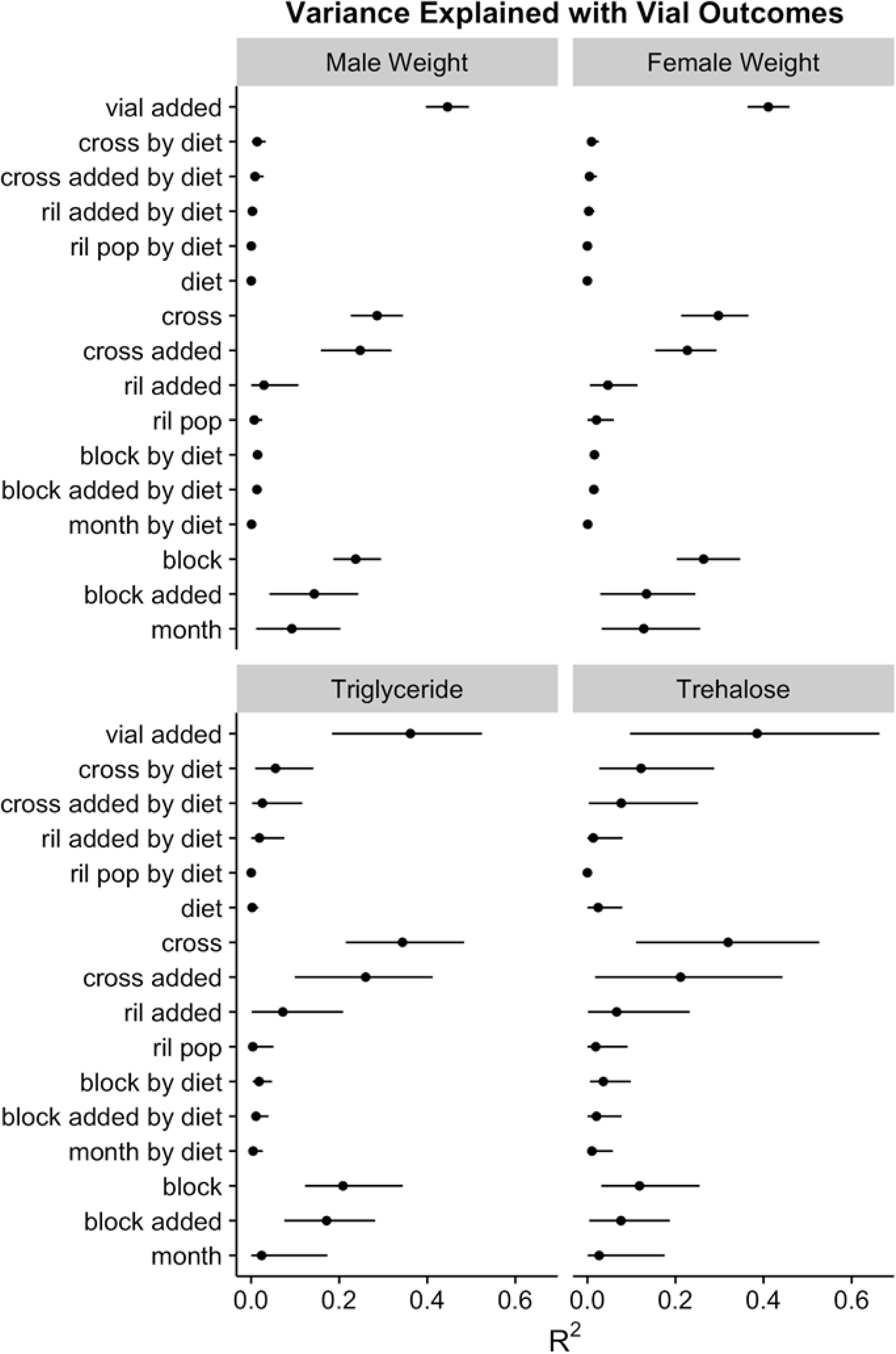
Variance explained by the null model for all phenotypes. The vial variance explained by each source of variation in the null model (month, block, RIL, cross, month by diet, block by diet, RIL by diet, cross by diet, and vial residuals) for male weight, female weight, triglyceride, and trehalose. The posterior mean and 95% credible interval are displayed above and raw values are in Table S6.

**Figure S2:**
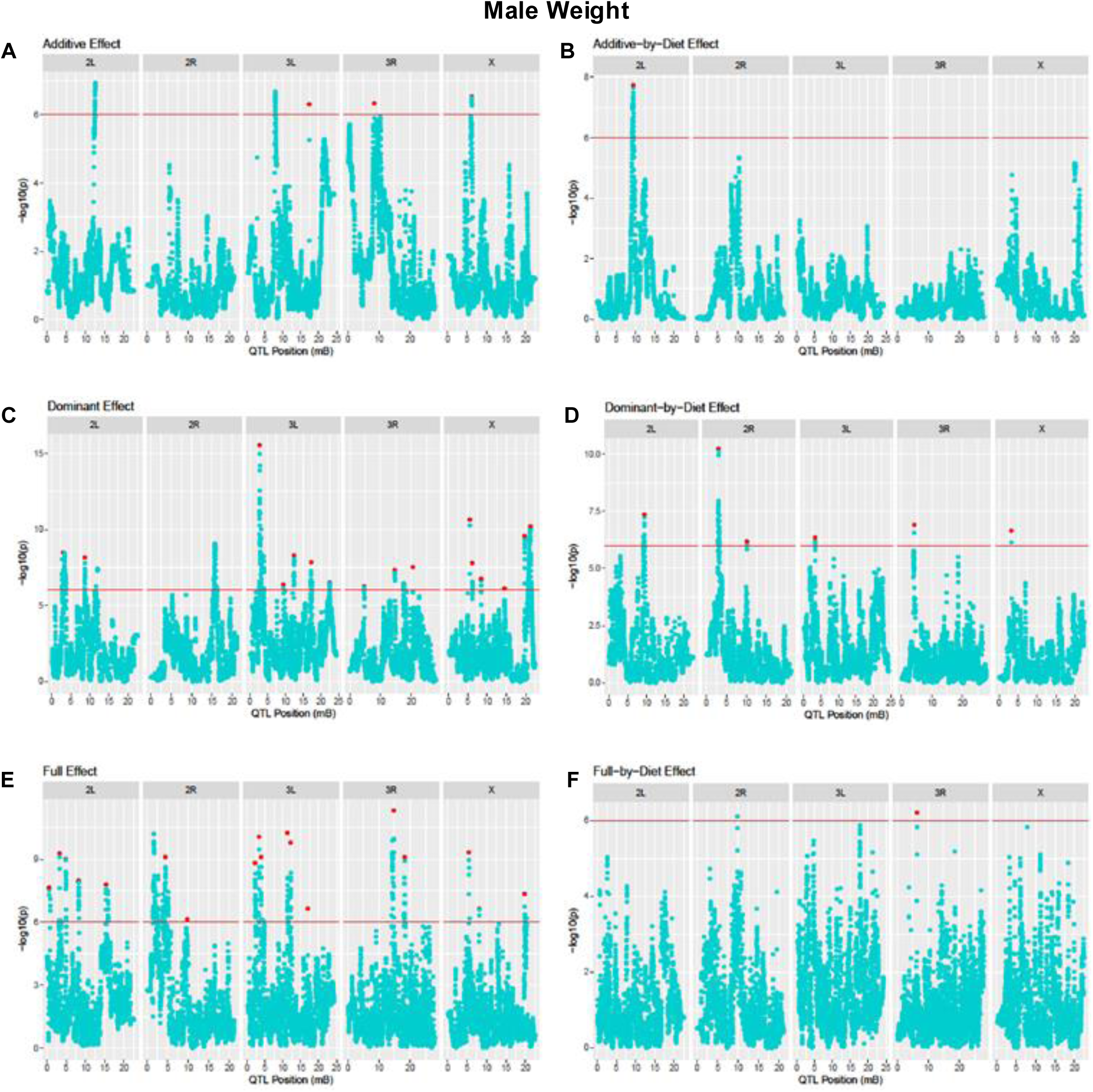
Non-epistatic genetic and genetic-by-diet effect Manhattan plots for the male weight phenotype. Manhattan plots showing the negative log p-value for the genetic effects (additive, A, dominant, C, and full, E, models) for the complete genome in the left-hand panels, while the right-hand panels indicate the genetic-by-diet effects (additive-by-diet, B, dominant-by-diet, D, and full-by-diet, F), for male weight phenotype. Red lines indicate the significance threshold and local maxima peaks are indicated by red dots. Data available in Tables S7, S8.

**Figure S3:**
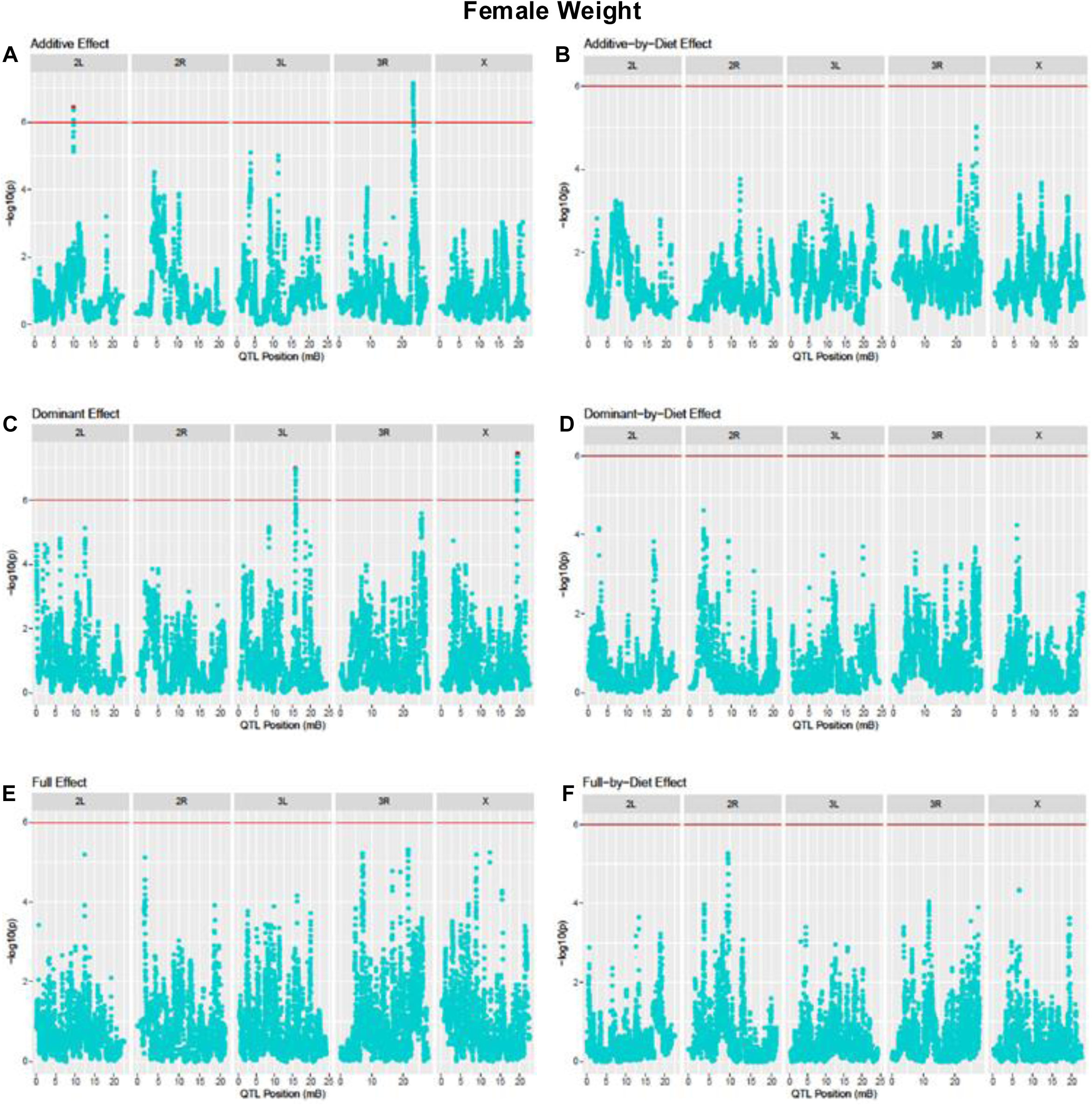
Non-epistatic genetic and genetic-by-diet effect Manhattan plots for the female weight phenotype. Manhattan plots showing the negative log p-value for the genetic effects (additive, A, dominant, C, and full, E, models) for the complete genome in the left-hand panels, while the right-hand panels indicate the genetic-by-diet effects (additive-by-diet, B, dominant-by-diet, D, and full-by-diet, F), for female weight phenotype. Red lines indicate the significance threshold and local maxima peaks are indicated by red dots. Data available in Tables S7, S9.

**Figure S4:**
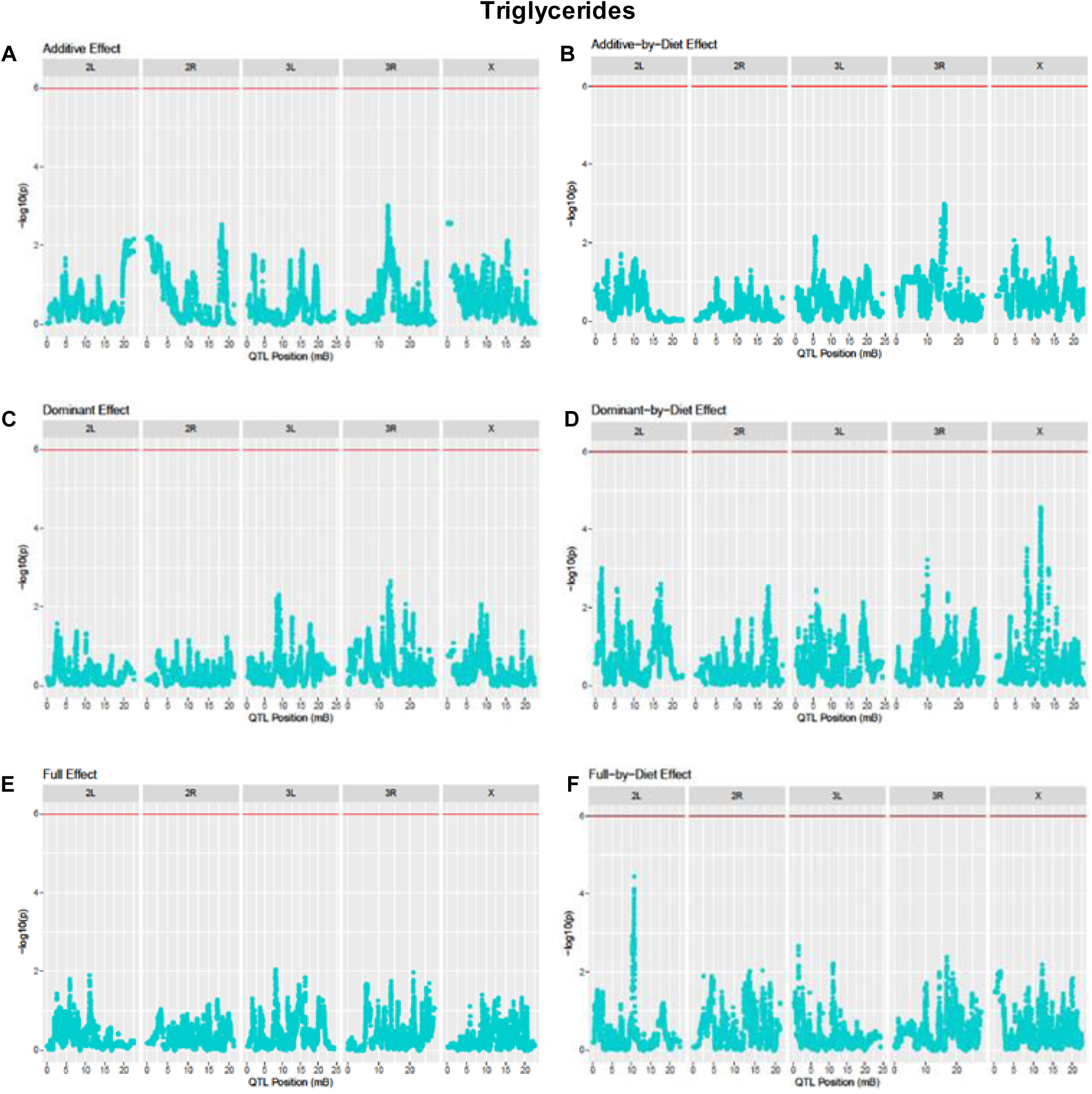
Non-epistatic genetic and genetic-by-diet effect Manhattan plots for the triglyceride phenotype. Manhattan plot showing the negative log p-value for the genetic effects (additive, A, dominant, C, and full, E, models) for the complete genome in the left-hand panels, while the right-hand panels indicate the genetic-by-diet effects (additive-by-diet, B, dominant-by-diet, D, and full-by-diet, F), for triglyceride phenotype. Red lines indicate the significance threshold and local maxima peaks are indicated by red dots. Data available in Tables S7, S10.

**Figure S5:**
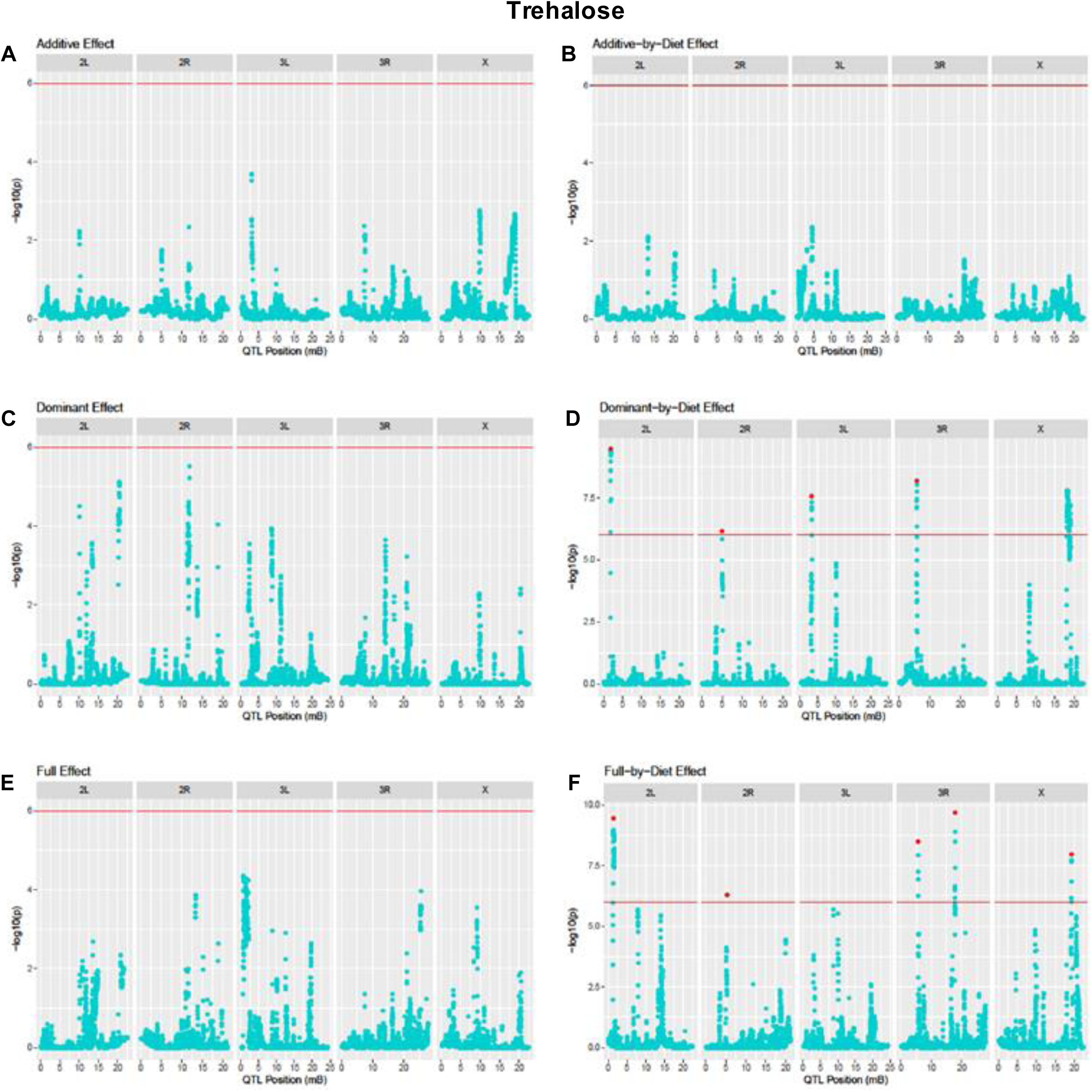
Non-epistatic genetic and genetic-by-diet effect Manhattan plots for the trehalose phenotypes. Manhattan plot showing the negative log p-value for the genetic effects (additive, A, dominant, C, and full, E, models) for the complete genome in the left-hand panels, while the right-hand panels indicate the genetic-by-diet effects (additive-by-diet, B, dominant-by-diet, D, and full-by-diet, F), for trehalose phenotype. Red lines indicate the significance threshold and local maxima peaks are indicated by red dots. Data available in Tables S7, S11.

**Figure S6:**
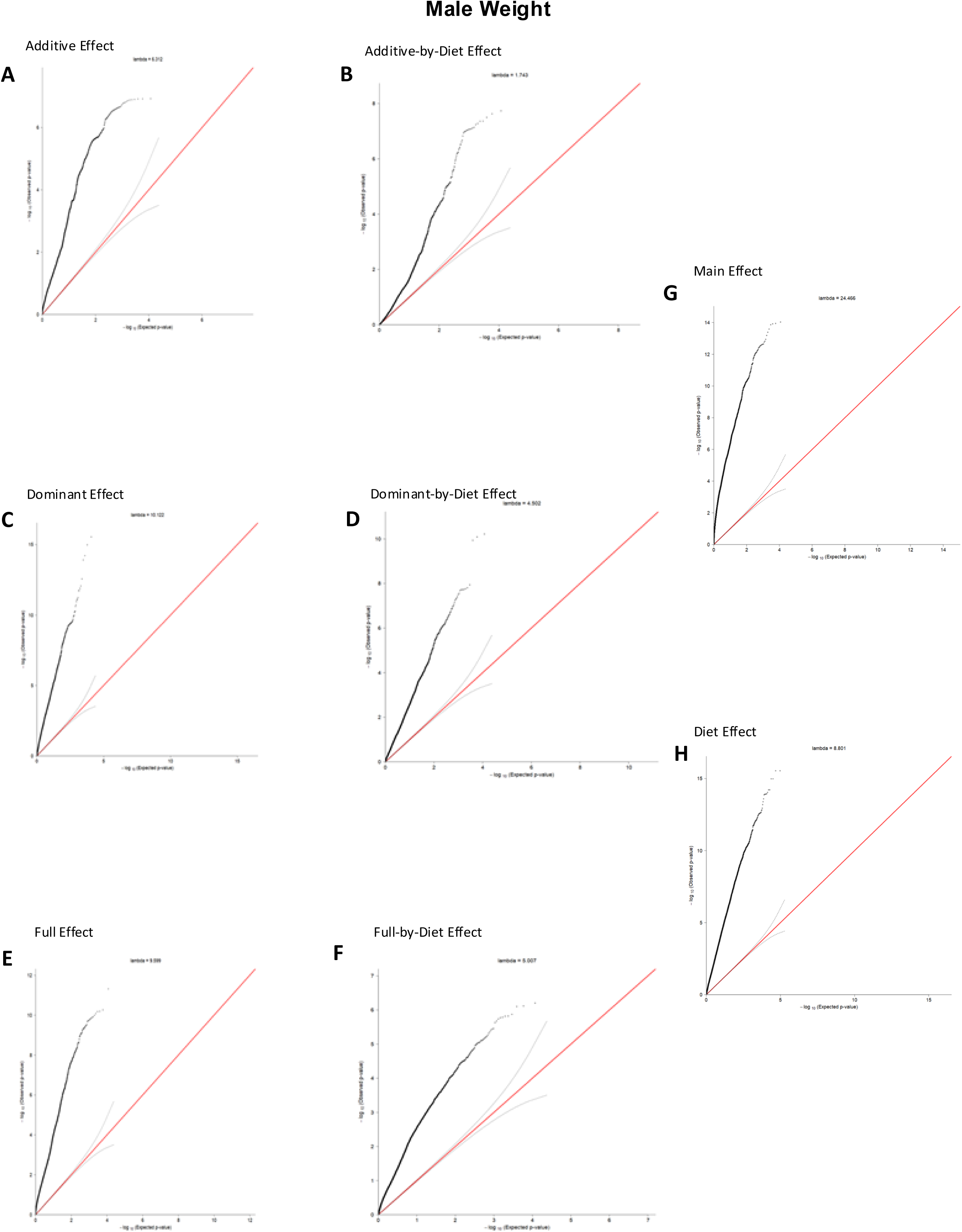
Non-epistatic genetic and genetic-by-diet effect QQ plots for the male weight phenotypes. Quantile-quantile plot showing the observed p-value (y-axis) compared to a standard uniform distribution (x-axis) for the additive (A), dominant (C), full (E), main (G), and diet (H) models.

**Figure S7:**
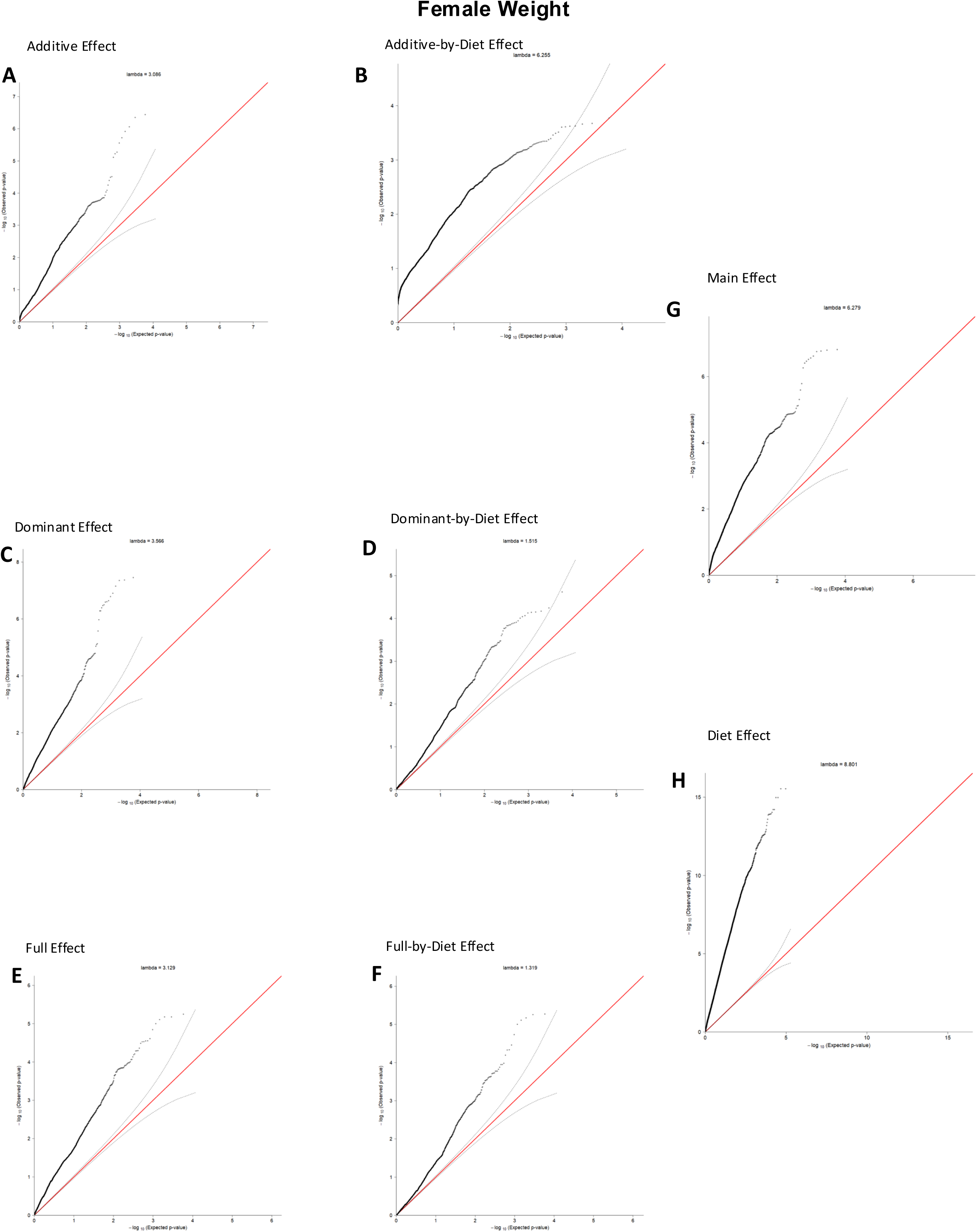
Non-epistatic genetic and genetic-by-diet effect QQ plots for the female weight phenotypes. Quantile-quantile plot showing the observed p-value (y-axis) compared to a standard uniform distribution (x-axis) for the additive (A), dominant (C), full (E), main (G), and diet (H) models.

**Figure S8:**
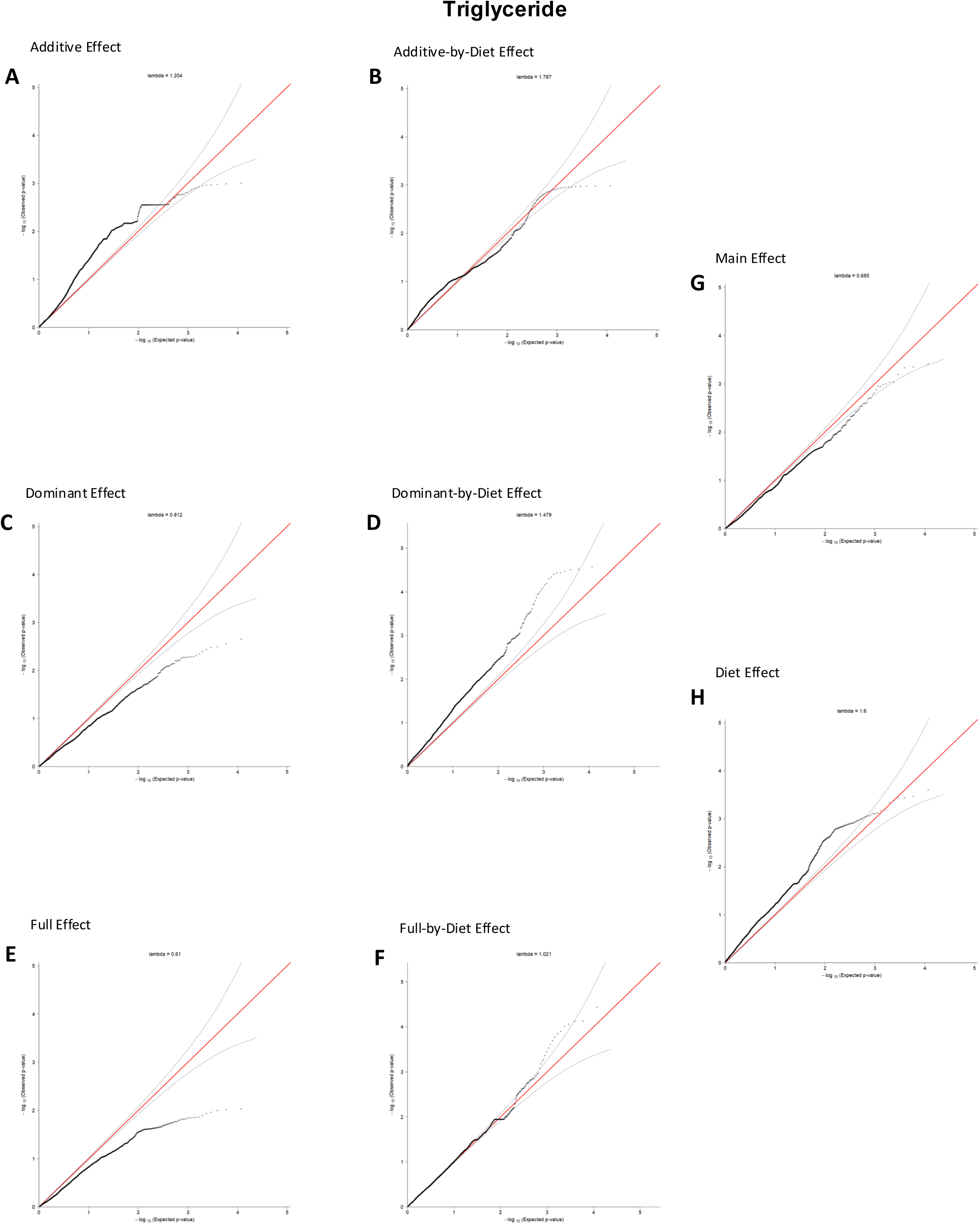
Non-epistatic genetic and genetic-by-diet effect QQ plots for the triglycerides phenotypes. Quantile-quantile plot showing the observed p-value (y-axis) compared to a standard uniform distribution (x-axis) for the additive (A), dominant (C), full (E), main (G), and diet (H) models.

**Figure S9:**
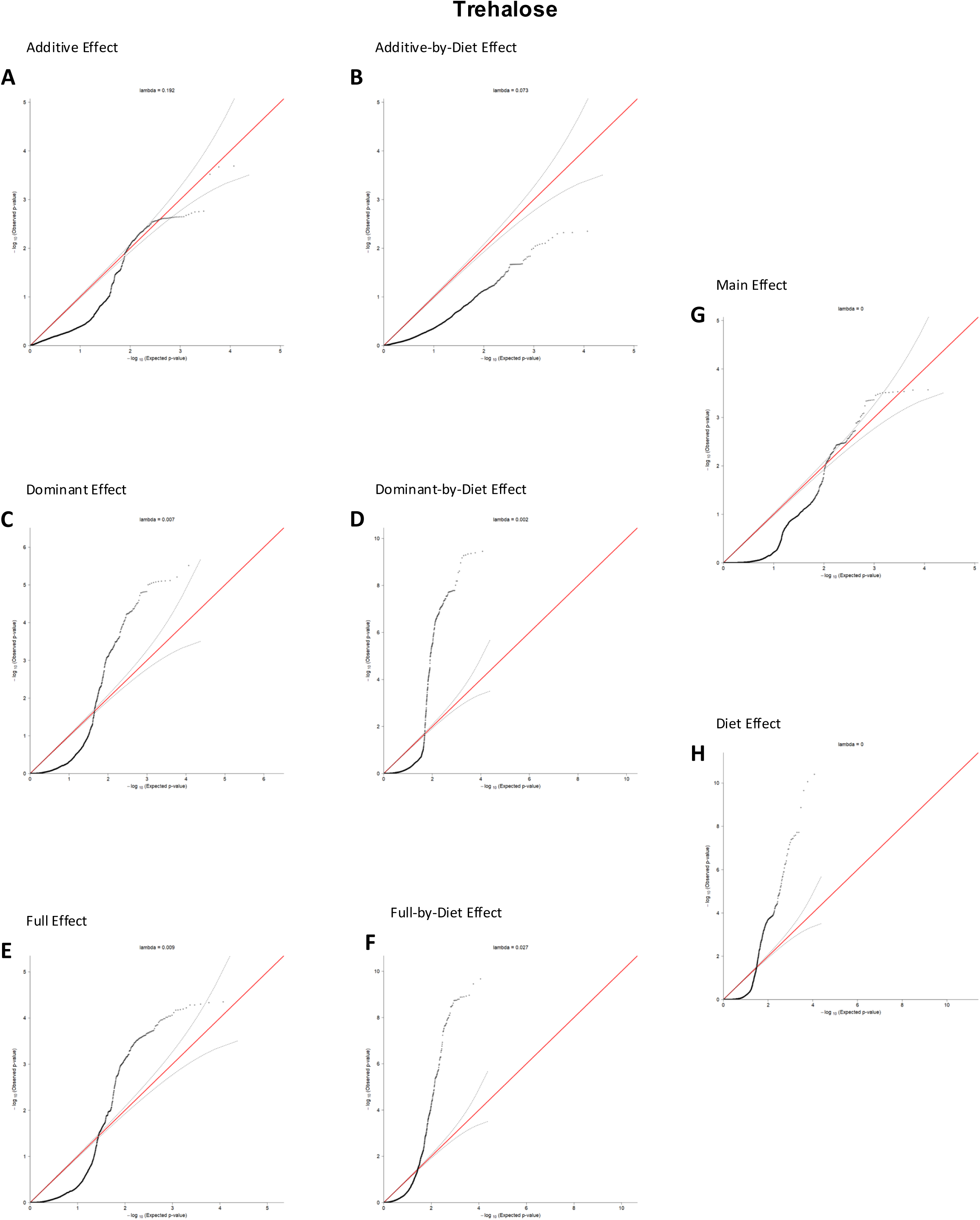
Non-epistatic genetic and genetic-by-diet effect QQ plots for the trehalose phenotypes. Quantile-quantile plot showing the observed p-value (y-axis) compared to a standard uniform distribution (x-axis) for the additive (A), dominant (C), full (E), main (G), and diet (H) models.

**Figure S10:**
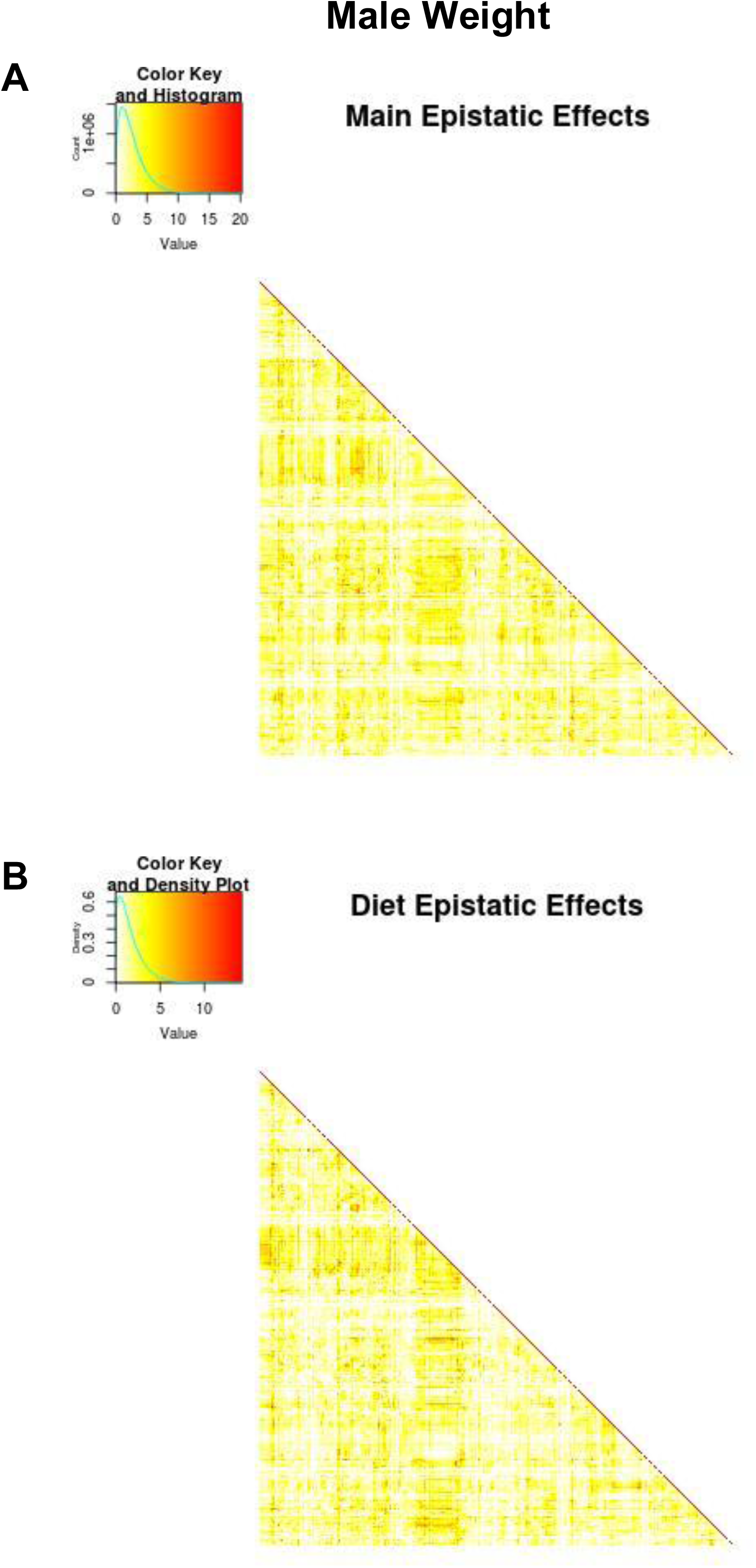
Male weight epistatic effect heat maps. Heat maps of p-values from main epistatic (A) and diet epistatic (B) effects for the male weight phenotype. All pairwise epistatic interactions were tested and p-values are shown on the negative log p-value scale with its density shown in the top left of the heat map. Data in Table S15.

**Figure S11:**
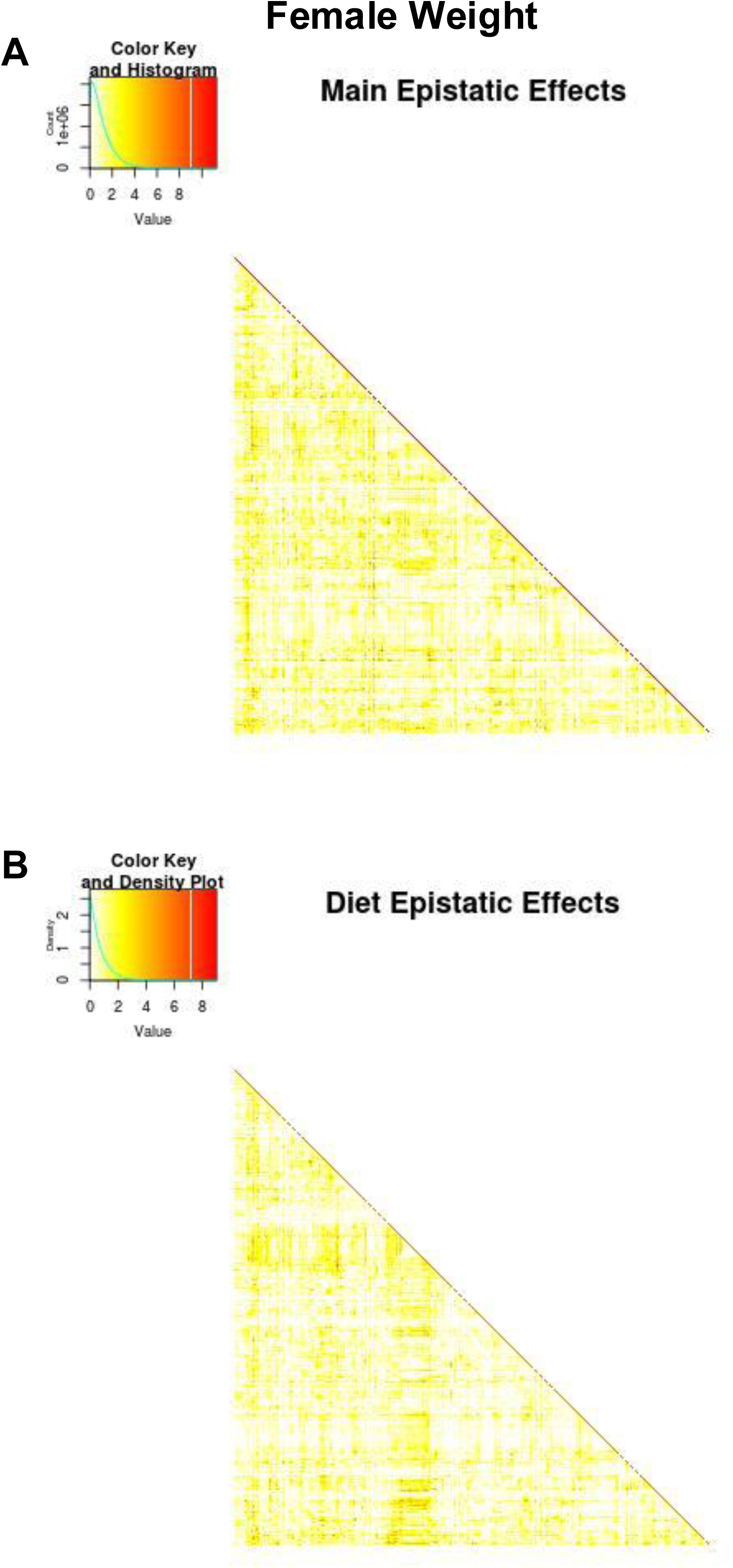
Female weight epistatic effect heat maps. Heat maps of p-values from main epistatic (A) and diet epistatic (B) effects for the female weight phenotype. All pairwise epistatic interactions were tested and p-values are shown on the negative log p-value scale with its density shown in the top left of the heat map. Data in Table S15.

**Figure S12.**
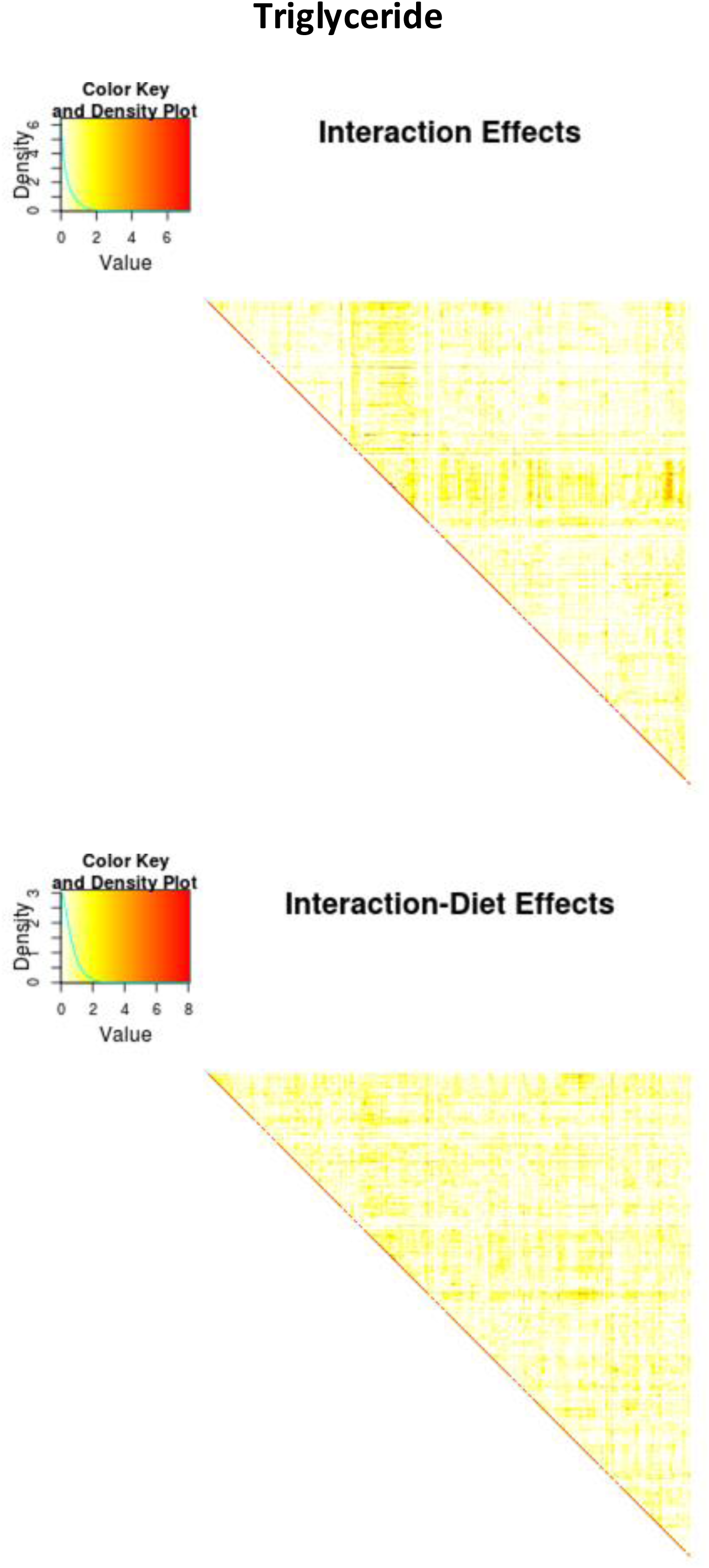
Triglyceride epistatic effect heat maps. Heat maps of p-values from main epistatic (A) and diet epistatic (B) effect for the triglyceride phenotype. All pairwise epistatic interactions were tested and p-values are shown on the negative log p-value scale with its density shown in the top left of the heat map. Data in Table S15.

**Figure S13:**
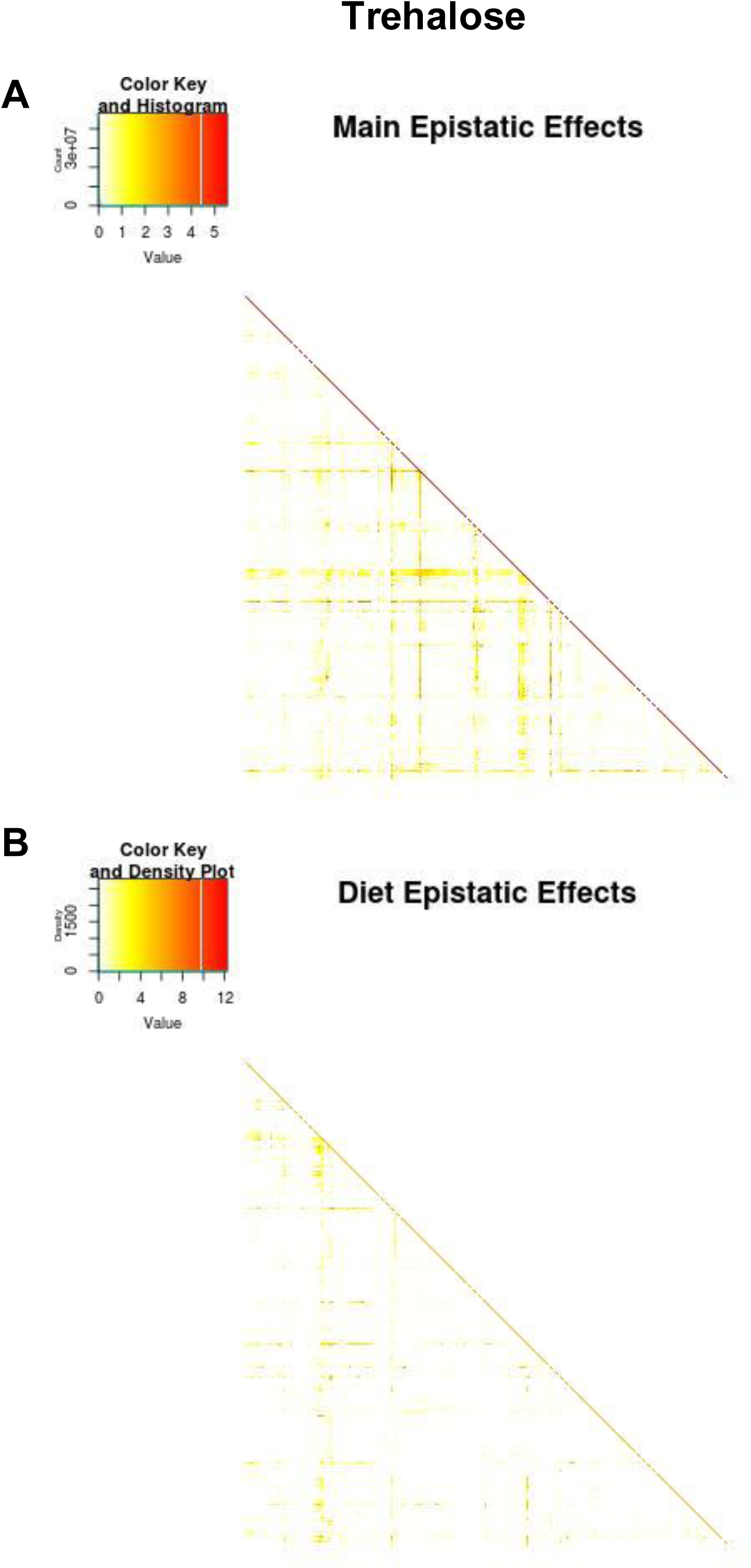
Trehalose epistatic effect heat maps. Heat maps of p-values from main epistatic (A) and diet epistatic (B) effect for the trehalose phenotype. All pairwise epistatic interactions were tested and p-values are shown on the negative log p-value scale with its density shown in the top left of the heat map. Data in Table S15.

**Figure S14:**
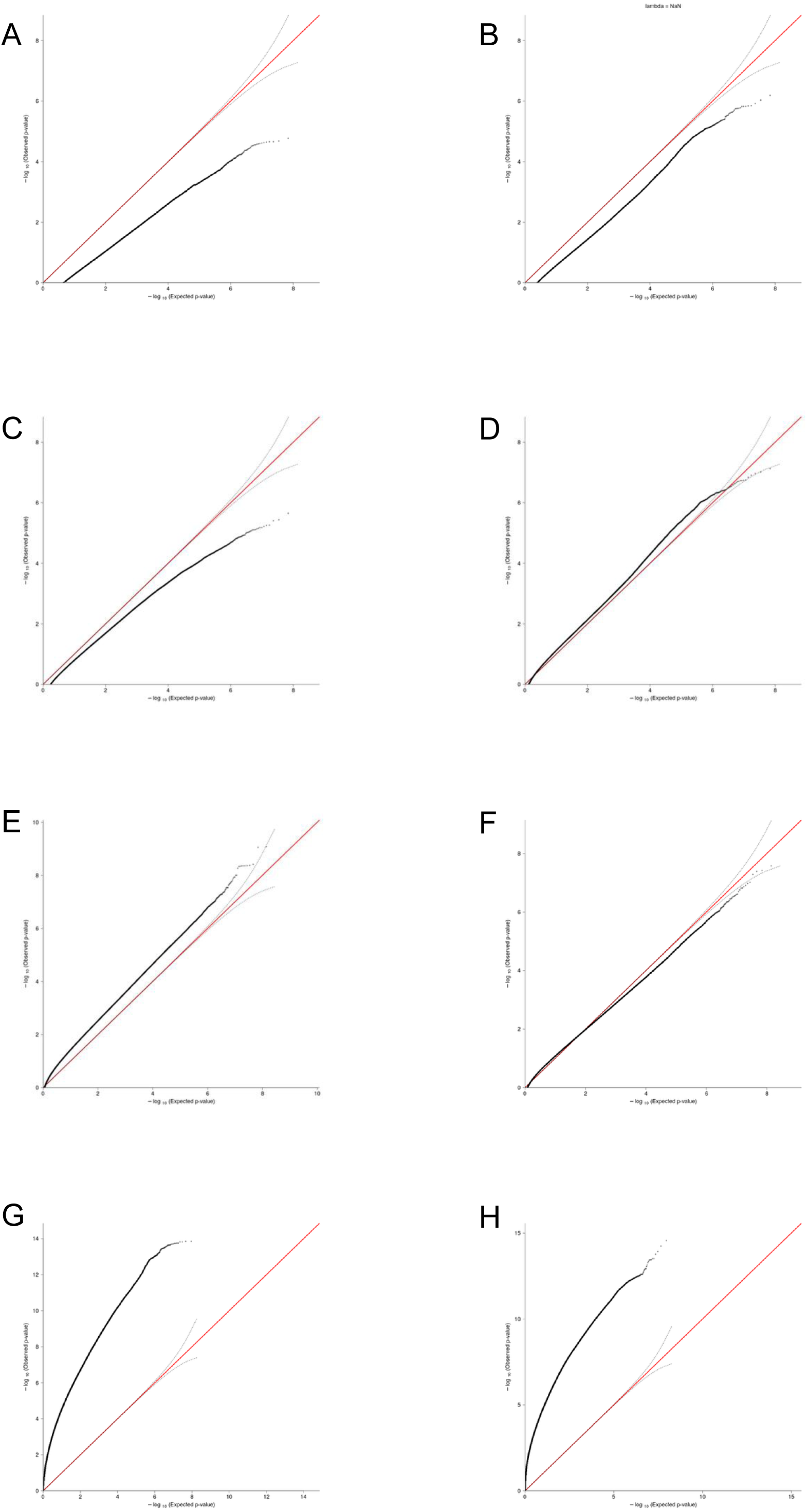
Epistatic genetic and genetic-by-diet effect QQ plots. Quantile-quantile plot showing the observed p-value (y-axis) compared to a standard uniform distribution (x-axis) for the male weight main (A), male weight diet (B), female weight main(C), female weight diet (D), triglyceride main (E), triglyceride diet (F), trehalose main (G), and trehalose diet (H) epistatic models.

**Figure S15:**
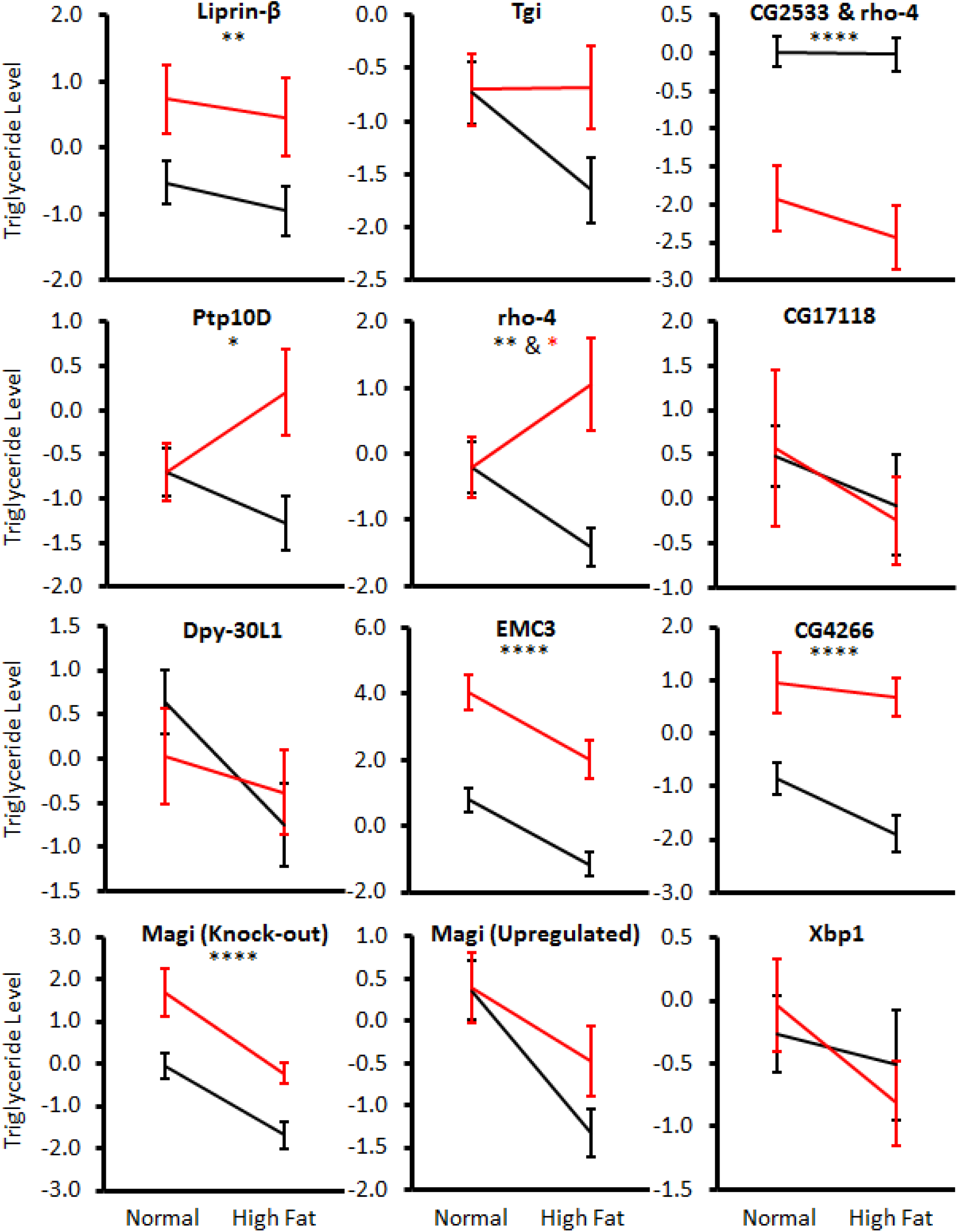
Candidate gene mutant triglycerides. Visualization of relationship between mutant (red) and control (black) triglycerides. All genes shown were significant for genotype (black stars) and one had a genotype-by-diet interaction (red star). FDR <0.3 used as multiple testing correction. 0.1 - 0.05 *, 0.05 – 0.01 **, 0.01 – 0.001 ***, <0.001 **** Data in Table S32.

**Figure S16:**
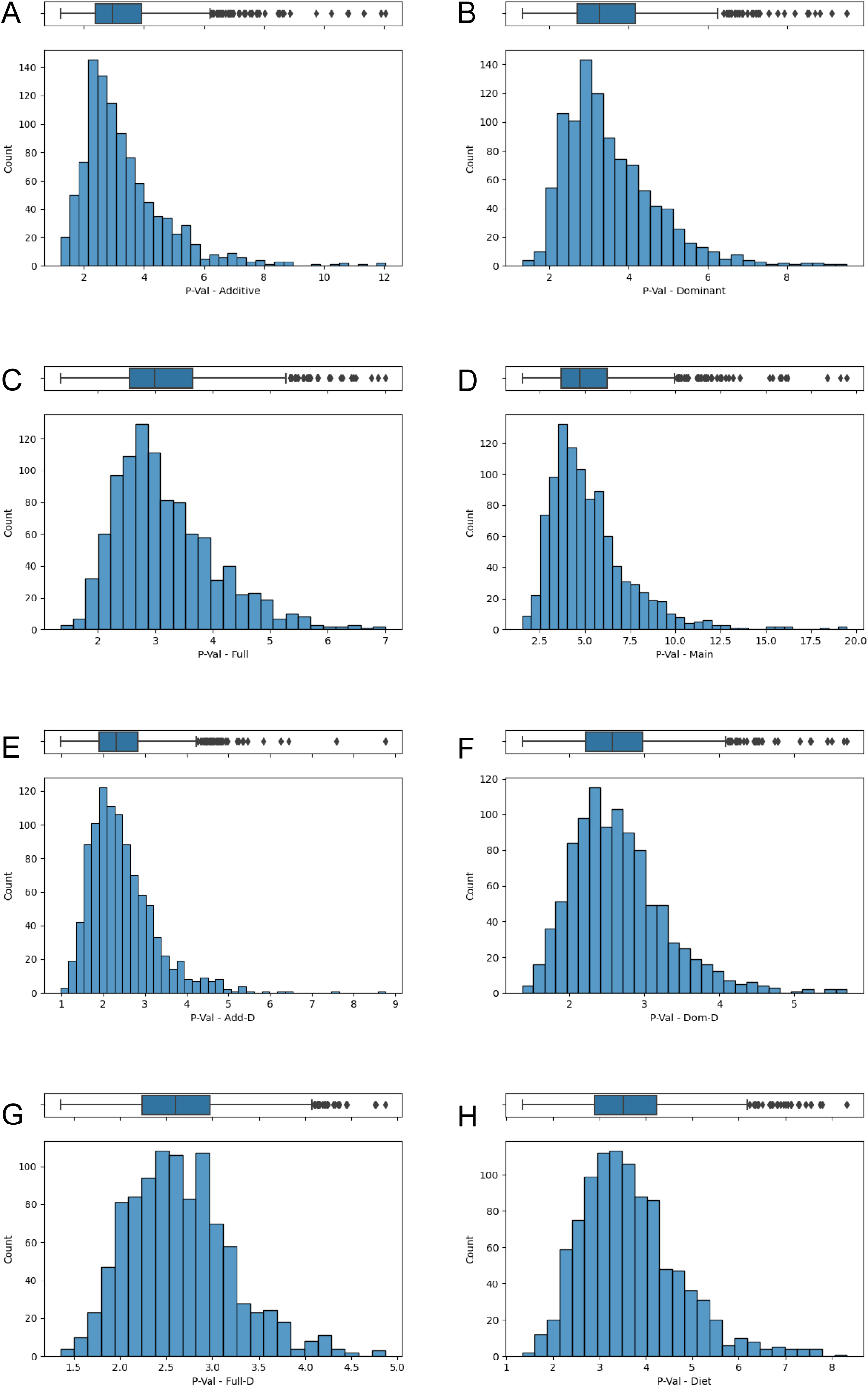
Non-epistatic male weight distributions of 5th percentile p-values (negative log p) from 1000 permutations. Permutation significance levels for each model (5th percentile of distribution of 5th percentiles) were (A) additive main: 6.19, (B) dominance main: 5.78, (C) full main: 4.92, (D) main: 9.52, (E) additive diet interaction: 4.03, (F) dominance diet interaction: 3.87, (G) full diet interaction: 3.71, and (H) diet interaction: 5.61. All data in Table S40.

Table S1: Overlap between non-epistatic candidate genes and other high-throughput studies.

Table S2: Overlap between epistatic candidate genes and other high-throughput studies.

Table S3: All results from functional testing of triglyceride candidate genes.

Table S4: Stocks used for functional testing of triglyceride candidate genes.

Table S5: Variance due to model effects for non-epistatic QTLs.

Table S6: Global variance explained by QTL model effects.

Table S7: All significant non-epistatic QTL peaks by phenotype and model.

Table S8: Complete non-epistatic mapping results for Male Weight.

Table S9: Complete non-epistatic mapping results for Female Weight.

Table S10: Complete non-epistatic mapping results for Triglyceride storage.

Table S11: Complete non-epistatic mapping results for Trehalose levels.

Table S12: All epistatic QTLs by phenotype and model.

Table S13: Interaction pairs between epistatic QTLs by phenotype and model.

Table S14: Overlap of epistatic clusters across phenotypes and models and associated candidate genes.

Table S15: Interactions among epistatic QTL clusters by phenotype and model

Table S16: All overlapping candidate genes for non-epistatic QTLs by phenotype and model

Table S17: Closest candidate genes within 20 kilobases of the peak if no overlapping gene.

Table S18: All candidate genes overlapping epistatic QTLs.

Table S19: Number of candidate genes overlapping epistatic QTL clusters.

Table S20: Epistatic candidate genes associated with two or more phenotypes.

Table S21: Ontology enrichment analysis for non-epistatic candidates in Drosophila.

Table S22: Ontology enrichment analysis for human orthologs of non-epistatic candidates.

Table S23: Ontology enrichment analysis for epistatic candidates in Drosophila.

Table S24: Ontology enrichment analysis for human orthologs of epistatic candidates.

Table S25: Ontology enrichment analysis for epistatic candidates in Drosophila overlapping two or more traits.

Table S26: Ontology enrichment analysis for human orthologs of epistatic candidates overlapping two or more traits.

Table S27: Ontology enrichment for non-epistatic candidate genes overlapping other studies.

Table S28: Ontology enrichment for epistatic candidate genes overlapping with other studies.

Table S29: Ontology enrichment for epistatic candidate genes overlapping with two or more other studies.

Table S30: Ontology enrichment for epistatic candidate genes overlapping with three or more other studies.

Table S31: Results of preliminary triglyceride QTL analysis used to selection genes for functional testing.

Table S32: Triglyceride candidate genes used in functional testing by associate QTL.

Table S33: Q-PCR results for testing candidate gene expression across diets.

Table S34: Complete mutant triglyceride phenotype data for candidate genes.

Table S35: Overlap between functionally triglyceride candidate genes and candidate genes for other phenotypes in this study.

Table S36: Number of epistatic cluster interactions versus cluster group size.

Table S37: Candidate genes found in both non-epistatic and epistatic mapping models.

Table S38: Ontology enrichment for candidate genes found in both non-epistatic and epistatic mapping models.

Table S39: Ontology enrichment for the 22 miR-312 associate candidate genes female weight epistatic QTLs.

Table 40: Non-epistatic male weight 5th percentile p-values (negative log p) from 1000 permutations for each of the eight models.

Table S41: Assembly position conversion between version 5 and version 6 of the *Drosophila melanogaster* genome assembly.

Table S42: Primers used for Q-PCR triglyceride candidate gene expression analysis.

Supplementary File S1: Supplementary methods

## REFERENCES

Alberti K. G. M. M., Zimmet P., Shaw J., 2006 Metabolic syndrome - a new world-wide definition. A Consensus Statement from the International Diabetes Federation. Diabet Med 23: 469–480.

Almén M. S., Nilsson E. K., Jacobsson J. A., Kalnina I., Klovins J., Fredriksson R., Schiöth H. B., 2014 Genome-wide analysis reveals DNA methylation markers that vary with both age and obesity. Gene 548: 61–67.

Arif S., Murat S., Almudi I., Nunes M. D. S., Bortolamiol-Becet D., McGregor N. S., Currie J. M. S., Hughes H., Ronshaugen M., Sucena É., Lai E. C., Schlötterer C., McGregor A. P., 2013 Evolution of mir-92a underlies natural morphological variation in Drosophila melanogaster. Curr Biol 23: 523–528.

Aybar M. J., Nieto M. A., Mayor R., 2003 Snail precedes slug in the genetic cascade required for the specification and migration of the Xenopus neural crest. Development 130: 483–494.

Barabási A.-L., Oltvai Z. N., 2004 Network biology: understanding the cell’s functional organization. Nat Rev Genet 5: 101–113.

Basciano H., Federico L., Adeli K., 2005 Fructose, insulin resistance, and metabolic dyslipidemia. Nutr Metab 2: 5.

Baumbach J., Hummel P., Bickmeyer I., Kowalczyk K. M., Frank M., Knorr K., Hildebrandt A., Riedel D., Jäckle H., Kühnlein R. P., 2014 A Drosophila in vivo screen identifies store-operated calcium entry as a key regulator of adiposity. Cell Metab 19: 331–343.

Benjamini Y., Hochberg Y., 1995 Controlling the False Discovery Rate: A Practical and Powerful Approach to Multiple Testing. J. R. Stat. Soc. Ser B (Methodol.) 57: 289–300.

Bi P., Kuang S., 2015 Notch signaling as a novel regulator of metabolism. Trends Endocrinol Metab 26: 248–255.

Bi P., Shan T., Liu W., Yue F., Yang X., Liang X.-R., Wang J., Li J., Carlesso N., Liu X., Kuang S., 2014 Inhibition of Notch signaling promotes browning of white adipose tissue and ameliorates obesity. Nat Med 20: 911–918.

Boutet A., De Frutos C. A., Maxwell P. H., Mayol M. J., Romero J., Nieto M. A., 2006 Snail activation disrupts tissue homeostasis and induces fibrosis in the adult kidney. EMBO J 25: 5603–5613.

Boyle E. A., Li Y. I., Pritchard J. K., 2017 An expanded view of complex traits: from polygenic to omnigenic. Cell 169: 1177–1186.

Bradford M. M., 1976 A rapid and sensitive method for the quantitation of microgram quantities of protein utilizing the principle of protein-dye binding. Anal Biochem 72: 248–254.

Bray M. S., Young M. E., 2007 Circadian rhythms in the development of obesity: potential role for the circadian clock within the adipocyte. Obes Rev 8: 169–181.

Broman K. W., Sen S., 2009 A Guide to QTL Mapping with R/qtl. Springer, New York.

Broughton S. J., Piper M. D. W., Ikeya T., Bass T. M., Jacobson J., Driege Y., Martinez P. Hafen E., Withers D. J., Leevers S. J., Partridge L., 2005 Longer lifespan, altered metabolism, and stress resistance in Drosophila from ablation of cells making insulin-like ligands. Proc Natl Acad Sci U.S.A. 102: 3105–3110.

Buniello A., MacArthur J. A. L., Cerezo M., Harris L. W., Hayhurst J., Malangone C., McMahon A., Morales J., Mountjoy E., Sollis E., Suveges D., Vrousgou O., Whetzel P. L., Amode R., Guillen J. A., Riat H. S., Trevanion S. J., Hall P., Junkins H., Flicek P., Burdett T., Hindorff L. A., Cunningham F., Parkinson H., The NHGRI-EBI GWAS Catalog of published genome-wide association studies, targeted arrays and summary statistics 2019 Nucleic Acids Res 47:D1005–D1012.

Chen A., Liu Y., Williams S. M., Morris N., Buchner D. A., 2017 Widespread epistasis regulates glucose homeostasis and gene expression (GS Barsh, Ed.). PLoS Genet 13: e1007025.

Chen Y.-W., Song S., Weng R., Verma P., Kugler J.-M., Buescher M., Rouam S., Cohen S. M., 2014 Systematic study of Drosophila microRNA functions using a collection of targeted knockout mutations. Dev Cell 31: 784–800.

Chopra V. S., Srinivasan A., Kumar R. P., Mishra K., Basquin D., Docquier M., Seum C., Pauli D., Mishra R. K., 2008 Transcriptional activation by GAGA factor is through its direct interaction with dmTAF3. Devel Biol 317: 660–670.

Clark A. G., Keith L. E., 1988 Variation among Extracted Lines of Drosophila-Melanogaster in Triacylglycerol and Carbohydrate Storage. Genetics 119: 595–607.

Clément K., Vaisse C., Manning B. S., Basdevant A., Guy-Grand B., Ruiz J., Silver K. D., Shuldiner A. R., Froguel P., Strosberg A. D., 1995 Genetic variation in the beta 3-adrenergic receptor and an increased capacity to gain weight in patients with morbid obesity. New Engl J Med 333: 352–354.

Corella D., Peloso G., Arnett D. K., Demissie S., Cupples L. A., Tucker K., Lai C. Q., Parnell L. D., Coltell O., Lee Y. C., Ordovas J. M., 2009 APOA2, dietary fat, and body mass index: replication of a gene-diet interaction in 3 independent populations. Arch Intern Med 169: 1897–1906.

Çiçek I. Ö., Karaca S., Brankatschk M., Eaton S., Urlaub H., Shcherbata H. R., 2016 Hedgehog Signaling Strength Is Orchestrated by the mir-310 Cluster of MicroRNAs in Response to Diet. Genetics 202: 1167–1183.

Crow J. F., 2010 On epistasis: why it is unimportant in polygenic directional selection. Philos Trans R Soc Lond B Biol Sci 365: 1241–1244.

Das U. N., Rao A. A., 2007 Gene expression profile in obesity and type 2 diabetes mellitus. Lipids Health Dis 6: 35.

De, R., Hu, T., Moore, J.H., Gilbert-Diamond, D.. 2015 Characterizing gene-gene interactions in a statistical epistasis network of twelve candidate genes for obesity. BioData Mining 8: 45.

De Luca M., Yi N. J., Allison D. B., Leips J., Ruden D. M., 2005 Mapping quantitative trait loci affecting variation in Drosophila triacylglycerol storage. Obes Res 13: 1596–1605.

Dzitoyeva S., Manev H., 2013 Reduction of Cellular Lipid Content by a Knockdown of Drosophila PDP1 γ and Mammalian Hepatic Leukemia Factor. J Lipids 2013: 297932–7.

Farooqi S., O’Rahilly S., 2006 Genetics of obesity in humans. Endocr Rev 27: 710–718.

Fetissov S. O., Meguid M. M., Sato T., Zhang L.-H., 2002 Expression of dopaminergic receptors in the hypothalamus of lean and obese Zucker rats and food intake. Am J Physiol Regul Integr Comp Physiol 283: R905–10.

Forsberg S., Bloom J., Sadhu M., Kruglyak L., Carlborg O., 2017. Accounting for genetic interactions improves modeling of individual quantitative trait phenotypes in yeast. Nat Genet 49: 497–503.

Garlapow M. E., Huang W., Yarboro M. T., Peterson K. R., Mackay T. F. C., 2015 Quantitative Genetics of Food Intake in Drosophila melanogaster. (DC Ko, Ed.). PLoS ONE 10: e0138129.

Garrido D., Rubin T., Poidevin M., Maroni B., Le Rouzic A., Parvy J.-P., Montagne J., 2015 Fatty acid synthase cooperates with glyoxalase 1 to protect against sugar toxicity. (R Kuhnlein, Ed.). PLoS Genet 11: e1004995.

Gelman A., Stern H., 2006 The Difference Between “Significant” and “Not Significant” is not Itself Statistically Significant. *Amer*. Stat., 60:328–331.

Ghezzi A., Zomeno M., Pietrzykowski A. Z., Atkinson N. S., 2016 Immediate-early alcohol-responsive miRNA expression in Drosophila. J Neurogenet 30: 195–204.

Gibson G., Reed L. K., 2008 Cryptic genetic variation. Curr Biol 18: R989–90.

Gjesing A. P., Sparsø T., Borch-Johnsen K., Jørgensen T., Pedersen O., Hansen T., Olsen N. V., 2009 No consistent effect of ADRB2 haplotypes on obesity, hypertension and quantitative traits of body fatness and blood pressure among 6,514 adult Danes. (AE Toland, Ed.). PLoS ONE 4: e7206.

Guan X. L., Cestra G., Shui G., Kuhrs A., Schittenhelm R. B., Hafen E., van der Goot F. G., Robinett C. C., Gatti M., Gonzalez-Gaitan M., Wenk M. R., 2013 Biochemical membrane lipidomics during Drosophila development. Dev Cell 24: 98–111.

Hill W. G., Goddard M. E., Visscher P. M., 2008 Data and theory point to mainly additive genetic variance for complex traits. PLoS Genet 4: e1000008.

Hirosumi J., Tuncman G., Chang L., Görgün C. Z., Uysal K. T., Maeda K., Karin M., Hotamisligil G. S., 2002 A central role for JNK in obesity and insulin resistance. Nature 420: 333–336.

Hori K., Sen A., Kirchhausen T., Artavanis-Tsakonas S., 2011 Synergy between the ESCRT-III complex and Deltex defines a ligand-independent Notch signal. J Cell Biol 195: 1005–1015.

Huang W., Mackay T. F. C., 2016 The Genetic Architecture of Quantitative Traits Cannot Be Inferred from Variance Component Analysis. PLoS Genet 12: e1006421

Huang W. W., Richards S. S., Carbone M. A. M., Zhu D. D., Anholt R. R. H. R., Ayroles J. F. J., Duncan L. L., Jordan K. W. K., Lawrence F. F., Magwire M. M. M., Warner C. B. C., Blankenburg K. K., Han Y. Y., Javaid M. M., Jayaseelan J. J., Jhangiani S. N. S., Muzny D. D., Ongeri F. F., Perales L. L., Wu Y.-Q. Y., Zhang Y. Y., Zou X. X., Stone E. A. E., Gibbs R. A. R., Mackay T. F. C. T., 2012 Epistasis dominates the genetic architecture of Drosophila quantitative traits. Proc Natl Acad Sci U.S.A. 109: 15553–15559.

Hulsegge G., Boer J. M., van der Beek A. J., Verschuren W. M., Sluijs I., Vermeulen R., Proper K. I., 2016 Shift workers have a similar diet quality but higher energy intake than day workers. Scand J Work Environ Health 42: 459–468.

Jeong H., Tombor B., Albert R., Oltvai Z. N., Barabási A. L., 2000 The large-scale organization of metabolic networks. Nature 407: 651 EP ––654.

Jocken J. W. E., Blaak E. E., Schiffelers S., Arner P., van Baak M. A., Saris W. H. M., 2007 Association of a beta-2 adrenoceptor (ADRB2) gene variant with a blunted in vivo lipolysis and fat oxidation. Int J Obes 31: 813–819.

Johnson P. M., Kenny P. J., 2010 Addiction-like reward dysfunction and compulsive eating in obese rats: Role for dopamine D2 receptors. Nat Neurosci 13: 635–641.

Jumbo-Lucioni P., Ayroles J. F., Chambers M. M., Jordan K. W., Leips J., Mackay T. F., De Luca M., 2010 Systems genetics analysis of body weight and energy metabolism traits in Drosophila melanogaster. BMC Genomics 11: 297.

Jumbo-Lucioni P., Bu S., Harbison S. T., Slaughter J. C., Mackay T. F., Moellering D. R., De Luca M., 2012 Nuclear genomic control of naturally occurring variation in mitochondrial function in Drosophila melanogaster. BMC Genomics 13: 659–659.

Kadowaki H., Yasuda K., Iwamoto K., Otabe S., Shimokawa K., Silver K., Walston J., Yoshinaga H., Kosaka K., Yamada N., Saito Y., Hagura R., Akanuma Y., Shuldiner A., Yazaki Y., Kadowaki T., 1995 A Mutation in the β3-Adrenergic Receptor Gene Is Associated with Obesity and Hyperinsulinemia in Japanese Subjects. Biochem Bioph Res Co 215: 555–560.

Kaur J., 2014 A comprehensive review on metabolic syndrome. Cardiol Res Pract 2014: 943162–21.

Kayali A. G., Eichhorn J., Haruta T., Morris A. J., Nelson J. G., Vollenweider P., Olefsky J. M., Webster N. J., 1998 Association of the insulin receptor with phospholipase C-gamma (PLCgamma) in 3T3-L1 adipocytes suggests a role for PLCgamma in metabolic signaling by insulin. J Biol Chem 273: 13808–13818.

Kilpeläinen T. O., Lakka T. A., Laaksonen D. E., Mager U., Salopuro T., Kubaszek A., Todorova B., Laukkanen O., Lindström J., Eriksson J. G., Hämäläinen H., Aunola S., Ilanne-Parikka P., Keinänen-Kiukaanniemi S., Tuomilehto J., Laakso M., Uusitupa M., Finnish Diabetes Prevention Study Group, 2008 Interaction of single nucleotide polymorphisms in ADRB2, ADRB3, TNF, IL6, IGF1R, LIPC, LEPR, and GHRL with physical activity on the risk of type 2 diabetes mellitus and changes in characteristics of the metabolic syndrome: The Finnish Diabetes Prevention Study. Metab Clin Exp 57: 428–436.

King E. G., Kislukhin G., Walters K. N., Long A. D., 2014 Using Drosophila melanogaster to identify chemotherapy toxicity genes. Genetics 198: 31–43.

King E. G., Macdonald S. J., Long A. D., 2012a Properties and Power of the Drosophila Synthetic Population Resource for the Routine Dissection of Complex Traits. Genetics 191: 935–949.

King E. G., Merkes C. M., McNeil C. L., Hoofer S. R., Sen S., Broman K. W., Long A. D., Macdonald S. J., 2012b Genetic dissection of a model complex trait using the Drosophila Synthetic Population Resource. Genome Res 22: 1558–1566.

Langin D., 2006 Adipose tissue lipolysis as a metabolic pathway to define pharmacological strategies against obesity and the metabolic syndrome. Pharmacol Res 53: 482–491.

Laposky A. D., Bass J., Kohsaka A., Turek F. W., 2008 Sleep and circadian rhythms: key components in the regulation of energy metabolism. Febs Letters 582: 142–151.

Lehner B., Tischler J., Fraser A. G., 2006 RNAi screens in Caenorhabditis elegans in a 96-well liquid format and their application to the systematic identification of genetic interactions. Nat. Protoc 1:1617–1620.

Li Y., Li S., Li R., Xu J., Jin P., Chen L., Ma F., 2017 Genome-wide miRNA screening reveals miR-310 family members negatively regulate the immune response in Drosophila melanogaster via co-targeting Drosomycin. Dev Comp Immunol 68: 34–45.

Lorenzo M., Teruel T., Hernandez R., Kayali A. G., Webster N. J. G., 2002 PLCgamma participates in insulin stimulation of glucose uptake through activation of PKCzeta in brown adipocytes. Exp Cell Res 278: 146–157.

Lu J., Fu Y., Kumar S., Shen Y., Zeng K., Xu A., Carthew R., Wu C.-I., 2008 Adaptive evolution of newly emerged micro-RNA genes in Drosophila. Mol Biol Evol 25: 929–938.

Lu X., Jain V. V., Finn P. W., Perkins D. L., 2007 Hubs in biological interaction networks exhibit low changes in expression in experimental asthma. In: *EMBO Press*, p. 98.

Mackay, T. Epistasis and quantitative traits: using model organisms to study gene–gene interactions. 2014 Nat Rev Genet 15: 22–33.

Mackay T. F. C. T., Richards S. S., Stone E. A. E., Barbadilla A. A., Ayroles J. F. J., Zhu D. D., Casillas S. S., Han Y. Y., Magwire M. M. M., Cridland J. M. J., Richardson M. F. M., Anholt R. R. H. R., Barrón M. M., Bess C. C., Blankenburg K. P. K., Carbone M. A. M., Castellano D. D., Chaboub L. L., Duncan L. L., Harris Z. Z., Javaid M. M., Jayaseelan J. C. J., Jhangiani S. N. S., Jordan K. W. K., Lara F. F., Lawrence F. F., Lee S. L. S., Librado P. P., Linheiro R. S. R., Lyman R. F. R., Mackey A. J. A., Munidasa M. M., Muzny D. M. D., Nazareth L. L., Newsham I. I., Perales L. L., Pu L.-L. L., Qu C. C., Ràmia M. M., Reid J. G. J., Rollmann S. M. S., Rozas J. J., Saada N. N., Turlapati L. L., Worley K. C. K., Wu Y.-Q. Y., Yamamoto A. A., Zhu Y. Y., Bergman C. M. C., Thornton K. R. K., Mittelman D. D., Gibbs R. A. R., 2012 The Drosophila melanogaster Genetic Reference Panel. Nature 482: 173–178.

Manichaikul A., Dupuis J., Sen S., Broman K. W., 2006 Poor performance of bootstrap confidence intervals for the location of a quantitative trait locus. Genetics 174: 481–489.

McLaren W., Pritchard B., Rios D., Chen Y., Flicek P., Cunningham F., 2010 Deriving the consequences of genomic variants with the Ensembl API and SNP Effect Predictor. Bioinformatics 26: 2069–2070.

Moreno-Navarrete J. M., Botas P., Valdés S., Ortega F. J., Delgado E., Vázquez-Martín A., Bassols J., Pardo G., Ricart W., Menéndez J. A., Fernández-Real J. M., 2009 Val1483Ile in FASN gene is linked to central obesity and insulin sensitivity in adult white men. Obesity (Silver Spring) 17: 1755–1761.

Murillo-Maldonado J. M., Zeineddine F. B., Stock R., Thackeray J., Riesgo-Escovar J. R., 2011 Insulin receptor-mediated signaling via phospholipase C-γ regulates growth and differentiation in Drosophila. (EMC Skoulakis, Ed.). PLoS ONE 6: e28067.

Mudunuri U., Che A., Yi, M., Stephens, R.M., 2009 bioDBnet: the biological database network. Bioinformatics 25: 555–556.

Yi N., Diament A., Chiu S., Kim K., Allison D. B., Fisler J. S., Warden C. H., 2004 Genetics 167: 399-409

Nieto M. A., 2002 The snail superfamily of zinc-finger transcription factors. Nat Rev Mol Cell Biol 3: 155–166.

Ng’oma E., Fidelis W., Middleton K.M., King E.G., 2019, The evolutionary potential of diet-dependent effects on lifespan and fecundity in a multi-parental population of Drosophila melanogaster. Heredity 122:582–594.

Ng’oma E., Williams-Simon P.A., Rahman A., King E.G., 2020, Diverse biological processes coordinate the transcriptional response to nutritional changes in a Drosophila melanogaster multiparent population. BMC Genomics 21:84

Ooi S. L., Pan X., Peyser B. D., Ye P., Meluh P. B., Yuan D. S., Irizarry R. A., Bader J. S., Spencer F. A., Boeke J. D., 2006 Global Synthetic-lethal analysis and yeast functional profiling. Trends Genet, 22: 56 – 63

O’Rahilly S., Farooqi I. S., 2006 Genetics of obesity. Philos T R Soc B 361: 1095–1105.

Phillips P. C., 2008 Epistasis--the essential role of gene interactions in the structure and evolution of genetic systems. Nature 9: 855–867.

Pospisilik J. A., Schramek D., Schnidar H., Cronin S. J. F., Nehme N. T., Zhang X., Knauf C., Cani P. D., Aumayr K., Todoric J., Bayer M., Haschemi A., Puviindran V., Tar K., Orthofer M., Neely G. G., Dietzl G., Manoukian A., Funovics M., Prager G., Wagner O., Ferrandon D., Aberger F., Hui C.-C., Esterbauer H., Penninger J. M., 2010 Drosophila genome-wide obesity screen reveals hedgehog as a determinant of brown versus white adipose cell fate. Cell 140: 148–160.

Reed L.K., Williams S., Springston M., Brown J., Freeman K., DesRoches C.E., Sokolowski M.B., Gibson G., 2010 Genotype-by-Diet Interactions Drive Metabolic Phenotype Variation in Drosophila melanogaster. Genetics 185:1009–1019

Reed L. K., Lee K., Zhang Z., Rashid L., Poe A., Hsieh B., Deighton N., Glassbrook N., Bodmer R., Gibson G., 2014 Systems Genomics of Metabolic Phenotypes in Wild-Type Drosophila melanogaster. Genetics. 197.2: 781–793

Reed L. K., Williams S., Springston M., Brown J., Freeman K., DesRoches C. E., Sokolowski M. B., Gibson G., 2010 Genotype-by-diet interactions drive metabolic phenotype variation in Drosophila melanogaster. Genetics 185: 1009–1019.

Reimand J., Arak T., Adler P., Kolberg L., Reisberg S., Peterson H., Vilo J., 2016 g:Profiler-a web server for functional interpretation of gene lists (2016 update). Nucleic Acids Res 44: W83–9.

Rockman M. V., 2012 The QTN program and the alleles that matter for evolution: all that’s gold does not glitter. Evolution 66: 1–17.

Ropers F., Derivery E., Hu H., Garshasbi M., Karbasiyan M., Herold M., Nürnberg G., Ullmann R., Gautreau A., Sperling K., Varon R., Rajab A., 2011 Identification of a novel candidate gene for non-syndromic autosomal recessive intellectual disability: the WASH complex member SWIP. Hum Mol Genet 20: 2585–2590.

Ross S. E., Hemati N., Longo K. A., Bennett C. N., Lucas P. C., Erickson R. L., MacDougald O. A., 2000 Inhibition of adipogenesis by Wnt signaling. Science 289: 950–953.

Rulifson E. J., Kim S. K., Nusse R., 2002 Ablation of insulin-producing neurons in flies: Growth and diabetic phenotypes. Science 296: 1118–1120.

Ryabinina O. P., Subbian E., Iordanov M. S., 2006 D-MEKK1, the Drosophila orthologue of mammalian MEKK4/MTK1, and Hemipterous/D-MKK7 mediate the activation of D-JNK by cadmium and arsenite in Schneider cells. BMC Cell Biol 7: 7.

Sanchez-Díaz I., Rosales-Bravo F., Reyes-Taboada J. L., Covarrubias A. A., Narvaez-Padilla V., Reynaud E., 2015 The Esg Gene Is Involved in Nicotine Sensitivity in Drosophila melanogaster. (G Roman, Ed.). PLoS ONE 10: e0133956.

Schleinitz D., Klöting N., Körner A., Berndt J., Reichenbächer M., Tönjes A., Ruschke K., Böttcher Y., Dietrich K., Enigk B., Filz M., Schön M. R., Jenkner J., Kiess W., Stumvoll M., Blüher M., Kovacs P., 2010 Effect of genetic variation in the human fatty acid synthase gene (FASN) on obesity and fat depot-specific mRNA expression. Obesity (Silver Spring) 18: 1218–1225.

Schneider M., Troost T., Grawe F., Martinez-Arias A., Klein T., 2013 Activation of Notch in lgd mutant cells requires the fusion of late endosomes with the lysosome. J Cell Sci 126: 645– 656.

Schoenherr J. A., Drennan J. M., Martinez J. S., Chikka M. R., Hall M. C., Chang H. C., Clemens J. C., 2012 Drosophila activated Cdc42 kinase has an anti-apoptotic function. (H Steller, Ed.). PLoS Genet 8: e1002725.

Scott Chialvo C. H., Che R., Reif D., Motsinger-Reif A., Reed L. K., 2016 Eigenvector metabolite analysis reveals dietary effects on the association among metabolite correlation patterns, gene expression, and phenotypes. Metabolomics 12: 710.

Smemo, S., Tena, J., Kim, KH., Gamazon E. R., Sakabe N. J., Gómez-Marín C., Aneas I., Credidio F. L., Sobreira D. R., Wasserman N. F., Lee J. H., Puviindran V., Tam D., Shen M., Son J. E., Vakili N.A., Sung H. K., Naranjo S., Acemel R. D., Manzanares M., Nagy A., Cox N. J., Hui C.C., Gomez-Skarmeta J. L., Nóbrega M. A. 2014 Obesity-associated variants within *FTO* form long-range functional connections with *IRX3*. Nature 507: 371– 375.

St Pierre S. E., Ponting L., Stefancsik R., McQuilton P., FlyBase Consortium, 2014 FlyBase 102-advanced approaches to interrogating FlyBase. Nucleic Acids Re 42: D780–8.

Stanley P.D., Ng’oma E., O’Day S., King E.G., Genetic Dissection of Nutrition-Induced Plasticity in Insulin/Insulin-Like Growth Factor Signaling and Median Life Span in a Drosophila Multiparent Population. Genetics 206: 587–602.

Swarup S., Harbison S. T., Hahn L. E., Morozova T. V., Yamamoto A., Mackay T. F. C., Anholt R. R. H., 2012 Extensive epistasis for olfactory behaviour, sleep and waking activity in Drosophila melanogaster. Genet Res 94: 9–20.

Tan A. L. Y., Forbes J. M., Cooper M. E., 2007 AGE, RAGE, and ROS in diabetic nephropathy. Semin Nephrol 27: 130–143.

Tang Y., Feinberg T., Keller E. T., Li X.-Y., Weiss S. J., 2016 Snail/Slug binding interactions with YAP/TAZ control skeletal stem cell self-renewal and differentiation. Nat Cell Biol 18: 917–929.

Teleman A. A., Maitra S., Cohen S. M., 2006 Drosophila lacking microRNA miR-278 are defective in energy homeostasis. Gene Dev 20: 417–422.

Thankos P. K., Michaelides M., Umegaki H., Volkow N. D., 2008 D2R DNA Transfer Into the Nucleus Accumbens Attenuates Cocaine Self-Administration in Rats. *Synapse (New York*, N.Y.) 62: 481–486.

Trinh I., Boulianne G. L., 2013 Modeling obesity and its associated disorders in Drosophila. Physiology (Bethesda) 28: 117–124.

Tsurudome K., Tsang K., Liao E. H., Ball R., Penney J., Yang J.-S., Elazzouzi F., He T., Chishti A., Lnenicka G., Lai E. C., Haghighi A. P., 2010 The Drosophila miR-310 cluster negatively regulates synaptic strength at the neuromuscular junction. Neuron 68: 879–893.

Tyler A. L., Donahue L. R., Churchill G.A., Carter G. W., 2016 Weak epistasis generally stabilizes phenotypes in a mouse intercross. PLoS Genet. 12, e1005805 (2016).

Ugrankar R., Berglund E., Akdemir F., Tran C., Kim M. S., Noh J., Schneider R., Ebert B., Graff J. M., 2015 Drosophila glucome screening identifies Ck1alpha as a regulator of mammalian glucose metabolism. Nat Commun 6: 7102.

Urban S., Lee J. R., Freeman M., 2002 A family of Rhomboid intramembrane proteases activates all Drosophila membrane-tethered EGF ligands. EMBO J. 21: 4277–4286.

USDA National Nutrient Database for Standard Reference, 2015 USDA National Nutrient Database for Standard Reference. https://data.nal.usda.gov/dataset/composition-foods-raw-processed-prepared-usda-national-nutrient-database-standard-referenc-6

Vickers S. P., Dourish C. T., 2004 Serotonin receptor ligands and the treatment of obesity. Curr Opin Investig Drugs 5: 377–388.

Voigt J.-P., Schade R., Fink H., Hörtnagl H., 2002 Role of 5-HT1A receptors in the control of food intake in obese Zucker rats of different ages. Pharmacol Biochem Behav 72: 403–409.

Vonesch S. C., Lamparter D., Mackay T. F. C., Bergmann S., Hafen E., 2016 Genome-Wide Analysis Reveals Novel Regulators of Growth in Drosophila melanogaster. (GS Barsh, Ed.). PLoS Genet 12: e1005616.

Wei W.-H., Hemani G., Gyenesei A., Vitart V., Navarro P., Hayward C., Cabrera C. P., Huffman J. E., Knott S. A., Hicks A. A., Rudan I., Pramstaller P. P., Wild S. H., Wilson J. F., Campbell H., Hastie N. D., Wright A. F., Haley C. S., 2012 Genome-wide analysis of epistasis in body mass index using multiple human populations. Eur J Hum Genet 20: 857– 862.

Williams S., Dew-Budd K., Davis K., Anderson J., Bishop R., Freeman K., Davis D., Bray K., Perkins L., Hubickey J., Reed L. K., 2015 Metabolomic and Gene Expression Profiles Exhibit Modular Genetic and Dietary Structure Linking Metabolic Syndrome Phenotypes in Drosophila. G3 (Bethesda) 5: 2817–2829.

Xu P., Vernooy S. Y., Guo M., Hay B. A., 2003 The Drosophila microRNA Mir-14 suppresses cell death and is required for normal fat metabolism. Curr Biol 13: 790–795.

Xu X., Gopalacharyulu P., Seppänen-Laakso T., Ruskeepää A.-L., Aye C. C., Carson B. P., Mora S., Orešič M., Teleman A. A., 2012 Insulin signaling regulates fatty acid catabolism at the level of CoA activation. (E Rulifson, Ed.). PLoS Genet 8: e1002478.

Yan S. F., Ramasamy R., Schmidt A. M., 2009 Receptor for AGE (RAGE) and its ligands-cast into leading roles in diabetes and the inflammatory response. J Mol Med 87: 235–247.

Yatsenko A. S., Marrone A. K., Shcherbata H. R., 2014 miRNA-based buffering of the cobblestone-lissencephaly-associated extracellular matrix receptor dystroglycan via its alternative 3’-UTR. Nat Commun 5: 4906.

Zhao X., Feng D., Wang Q., Abdulla A., Xie X.-J., Zhou J., Sun Y., Yang E. S., Liu L.-P., Vaitheesvaran B., Bridges L., Kurland I. J., Strich R., Ni J.-Q., Wang C., Ericsson J., Pessin J. E., Ji J.-Y., Yang F., 2012 Regulation of lipogenesis by cyclin-dependent kinase 8-mediated control of SREBP-1. J Clin Invest 122: 2417–2427.

Zhou S., Mackay T. F. C., Anholt R. R. H., 2014 Transcriptional and epigenetic responses to mating and aging in Drosophila melanogaster. BMC Genomics 15: 927.

